# Regulation of a bacterial histidine kinase by a phase separating scaffolding protein

**DOI:** 10.1101/2021.07.20.452843

**Authors:** Chao Zhang, Wei Zhao, Samuel W. Duvall, Kimberly A. Kowallis, W. Seth Childers

**Affiliations:** Department of Chemistry, University of Pittsburgh, Pittsburgh, PA 15260, USA

**Keywords:** intrinsically disordered region (IDR), biomolecular condensate, histidine kinase, scaffold, *Caulobacter crescentus*, PAS domain

## Abstract

Scaffolding proteins customize the response of signaling networks to support cell development and behaviors. We investigated how the bacterial scaffolding protein PodJ regulates the histidine kinase PleC involved in the asymmetric cell division of *Caulobacter crescentus*. We reconstituted the PleC-PodJ signaling complex through both heterologous expression in *E. coli* and *in vitro* studies. *In vitro* PodJ phase separates as a biomolecular condensate that recruits and inhibits PleC kinase activity. By constructing an *in vivo* PleC-CcaS chimeric histidine kinase reporter assay, we have demonstrated how PodJ leverages its intrinsically disordered region (IDR) to bind and regulate PleC-CcaS signaling. Moreover, we observed that full-length PodJ_L_ regulates PleC-CcaS signaling, while a truncated PodJ_s_ could not regulate signaling activity. These results support a model where PodJ biomolecular condensate formation regulates the localization and activity of the cell fate determining kinase PleC.

## Introduction

Eukaryotic scaffolds add layers of regulation upon signaling pathways that include allosteric mechanisms^1^, feedback-regulation^2^, and phase separation into distinct compartments^3^. In comparison, the mechanism of how scaffolding proteins impact bacterial signaling pathways has been less studied. Three scaffolds play roles in regulating the subcellular position of histidine kinases involved in the asymmetric cell division of the gram-negative bacteria *Caulobacter crescentus*: PopZ-CckA^4^, SpmX-DivJ^5^, and PodJ-PleC^6,7^. PopZ^8^ and SpmX^9^ phase separate as a biomolecular condensate that recruits distinct clients, including the histidine kinase DivJ and CckA, and the pseudokinase DivL at the old cell pole^4^. At the opposite new cell pole, PodJ serves as a scaffold that sequesters four distinct clients directly or indirectly: PleC, PopA, DivL and CpaE^7^. In contrast to SpmX and PopZ, much less is known about how PodJ organizes biochemical events at the new cell pole. Therefore, in this study, we investigated how the scaffolding protein PodJ organizes and regulates the histidine kinase PleC.

Coordination of *C. crescentus* replication, cell growth, and division requires the bifunctional cell-cycle kinase CckA to undergo a kinase-to-phosphatase switch during each cell cycle. The activity changes of CckA are facilitated by the bifunctional histidine kinase PleC that also oscillates between kinase and phosphatase activity states^10,11^(Fig. 1A). During the swarmer-to-stalk transition, PleC phosphorylates its cognate response regulators PleD and DivK^12,13^. In turn, these phosphorylation events activate CckA’s phosphatase activity^10,14,15^ that inhibits the CtrA pathway and allows the initiation of chromosome replication. After the initiation of replication, PleC switches to its phosphatase state, leading to CtrA pathway activation^14,16^. Therefore, PleC’s cell-cycle mediated switch between kinase and phosphatase is crucial for cell development.

**Figure 1:**
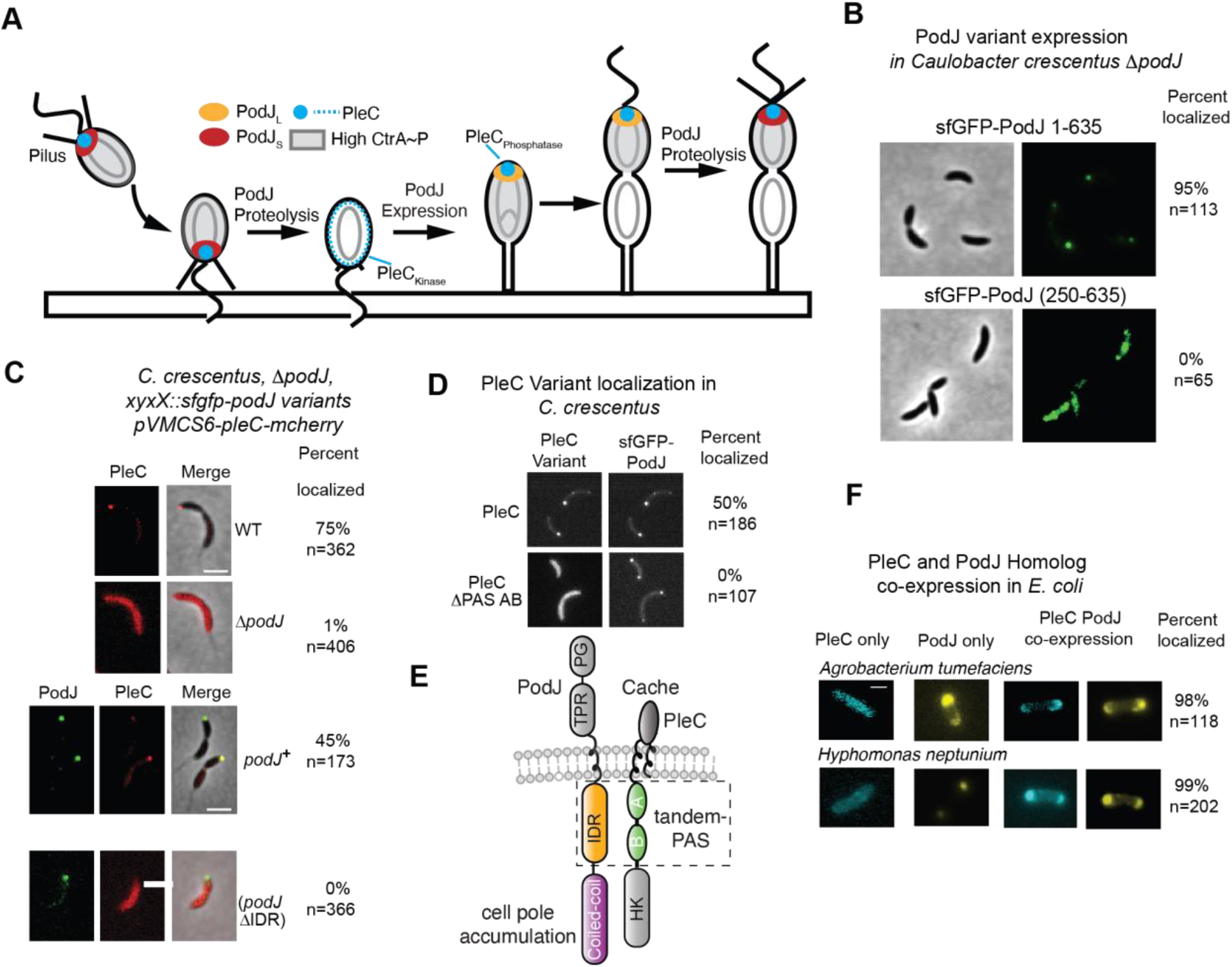
Identification of PodJ and PleC domains critical for co-localization at the cell pole. (A) The localization pattern of the PodJ-PleC signaling complex through the *Caulobacter crescentus* cell cycle. PodJi_L_ (orange) expression leads to cell pole accumulation of the PodJ-PleC and upregulation of PleC (blue) phosphatase function. Proteolysis of PodJ results in a shortened form of PodJ (red) that retains cell pole accumulation but does not stimulate PleC phosphatase function. Subsequent proteolysis of PodJ liberates PleC from the cell pole (B) Expression of sfGFP-PodJ(250-635) and sfGFP-PodJ(1-635) as a sole copy in *podJ* deletion *C. crescentus* strain. (C) Expression of PleC-mCherry in wild-type *C. crescentus,* the *podJ* deletion strain, and the *podJ* deletion strain supplemented full-length sfGFP-PodJ or sfGFP-PodJ-ΔIDR and their localization analysis. Scale bar: 2 μm. Strains were cultured into mid-log phase and induced with 0.03% xylose for sfGFP-PodJ and 0.05 mM vanillic acid for PleC-mCherry for 5 h before imaging. (D) Localization of PleC-mCherry or PleCΔPAS-AB with sfGFP-PodJ in the *C. crescentus podJ-pleC* deletion strain. Strains were cultured into the midlog phase and induced with 0.5 mM vanillic acid for PleC-mCherry for 3 h before imaging. Scale bar: 2 μm. For each image, the percent co-localized and the number of cells analyzed are reported. (E) Annotation of PodJ and PleC functions indicates an IDR-PAS protein-protein interaction and that PodJ’s N-terminal coiled coils are critical for cell pole accumulation. (F) YFP-PodJ and PleC-CFP pairs from other alpha-proteobacteria species co-localize at the cell poles when co-expressed in *E. coli.* YFP-PodJ was induced with 0.5 mM IPTG, and PleC-mCherry was induced with 1mM arabinose for 2 hours. Scale bar: 2 μm.

Several factors contribute to PleC’s kinase to phosphatase switch. One factor is that unphosphorylated DivK allosterically stimulates the kinase activity of PleC^17^. Supporting this model, it was shown the addition of unphosphorylatable DivK mutant, DivK D53N, stimulated PleC autophosphorylation *in vitro*^17^. Second, PleC signaling activity is regulated by pilus retraction upon surface contact^18^. In this model, pilus retraction leads to the accumulation of PilA monomers in the periplasm interacting with PleC^18^. This PilA-PleC interaction leads to an increase in cellular cyclic-di-GMP levels mediated by diguanylate cyclase PleD^18^. A third factor that correlates with PleC’s activity is its subcellular localization pattern (Fig. 1A). When PleC is localized at the new cell pole, it functions as a phosphatase^19^. In contrast, when PleC is released from the cell pole in the swarmer-to-stalk transition, PleC functions as a kinase^17^.

The cell pole localization pattern of PleC depends upon the scaffolding protein PodJ^6,20^. The PodJ scaffold spans the membrane and within the cytoplasm includes six coiled coils, an intrinsically disordered region (IDR) (Fig. S1A-S1B) featured by negatively and positively charged blocks at the N and C-terminal, respectively, and a transmembrane (TM) anchor. Within the periplasm, PodJ contains a tetratricopeptide repeat (TPR) domain and a peptidoglycan (PG)-binding domain. In *C. crescentus*, deletion of *podJ* results in delocalized PleC and downregulation of the CtrA pathway as indicated by a 47% reduced *pilA* promoter activity and reduced PilA accumulation in *podJ* mutants^7,20,21^. Moreover, PodJ variants lacking either cytoplasmic or periplasmic domains reduce *pilA* promoter activity^6^ and PleC cell pole localization^21^. Thus, both PodJ’s cytoplasmic and periplasmic domains contribute to the regulation of CtrA activity^21^.

Upon cell division, full-length PodJ_L_ (full length, 1-974) is proteolyzed into a shortened form PodJ_S_ (short form, 1-702),^22,23^ via a set of proteases^7,22,23^. Both PodJ_L_ and PodJ_S_ can support cell pole localization of PleC^21^. However, the proteolysis of PodJ correlates with a downregulation of the CtrA regulon^21^. Thus, PodJ may mediate changes to the CtrA regulatory pathway through PleC^6,20^. In addition, PodJ also regulates CtrA through another CtrA regulator, DivL^24^. Therefore, PodJ’s multifaceted impact upon CtrA regulators provides uncertainty about how PodJ regulates PleC’s function.

Here, we applied synthetic biology and *in vitro* biochemical approaches to interrogate if and how PodJ directly regulates PleC’s localization and activity. Our results indicate that the PodJ scaffold phase separates as a biomolecular condensate *in vitro* that recruits PleC and inhibits PleC kinase activity. Through a complementary synthetic biology strategy, we designed and built a PleC-CcaS chimeric histidine kinase reporter assay that identified domains necessary for the PodJ stimulation of PleC. Our results suggest a model in which PodJ phase separates as a biomolecular condensate that recruits and regulates PleC signaling activity.

## Results

### PodJ coiled-coils contribute to cell pole localization

To understand how PodJ impacts PleC’s function, we reconstituted the PodJ-PleC signaling complex in *E. coli*. Past studies have shown that heterologous expression of PodJ in *E. coli* resulted in cell pole accumulation^4^. Notably, the γ-proteobacterium *E. coli* is divergent from the alphaproteobacterium *C. crescentus* and does not contain any *C. crescentus* polarity protein homologs^25^. Therefore, *E. coli* has been used extensively as an orthologous system for testing *C. crescentus* protein-protein interactions^4,26^.

Previous studies have also shown that an N-terminal YFP fusion to PodJ does not disrupt its regulation of the CtrA pathway in *C. crescentus*^20^. Therefore, to reconstitute the PodJ-PleC complex, we heterologously expressed an N-terminal fluorescent fusion protein of PodJ(YFP-PodJ) in *E. coli* and determined its subcellular localization pattern. As shown in Fig. S2A, YFP-PodJ accumulated at the cell poles. We also found that PodJ variants lacking the periplasmic domains, transmembrane tether, IDR, or coiled-coil 4-6 (CC4-6) also accumulated at the cell poles. In contrast, PodJ variants lacking coiled-coil 1-3 (CC1-3) did not accumulate at the cell poles in *E. coli*(Fig. S2A).

In *C. crescentus*, Lawler *et al*. observed that both the cytoplasmic and the periplasmic domains alone of PodJ could accumulate at the cell poles^21^. We, therefore, interrogated the role of CC1-3 in the subcellular localization of PodJ’s cytoplasmic domains, YFP-PodJ(1-635). As in *E. coli*, we observed that YFP-PodJ(1-635) accumulated at the cell poles in *C. crescentus* (Fig. 1B). In contrast, we observed that sfGFP-PodJ(250-635) was diffuse or patchy when expressed as a sole copy in *C. crescentus* (Fig. 1B). This suggests that CC1-3 is critical for the ability of PodJ’s cytoplasmic domains to accumulate at the cell poles.

### PleC localizes to the cell pole via PodJ’s IDR

With an understanding of the domains that contribute to PodJ subcellular localization (Fig. 1B, S2A), we then interrogated if YFP-PodJ recruits PleC-mCherry to the cell poles in *E. coli.* The expression of PleC alone resulted in a diffuse localization pattern (Fig. S2A). In contrast, co-expression of PodJ and PleC resulted in co-localization at the cell poles in 98% of cells. To interrogate the interaction between YFP-PodJ and PleC-mCherry, we heterologously co-expressed full-length PleC-mCherry with a library of YFP-PodJ domain deletion variants in *E. coli* (Fig. S2A). Deletion of the periplasmic domains, the TM or CC4-6, did not affect the cell pole localization of YFP-PodJ or PleC-mCherry recruitment to the cell poles. In contrast, the YFP-PodJΔIDR variant was unable to recruit PleC-mCherry to the cell poles (Fig. S2A). This suggests that PodJ’s IDR may be a site of interaction with PleC-mCherry in *E. coli*.

We next tested if PodJ’s IDR was critical for PleC localization in *C. crescentus* (Fig.1C). Past work has shown that PleC-GFP localized at the new cell pole in pre-divisional cells^19^, and was dependent upon PodJ^21^. Consistent with these findings, we observed that in a *podJ* deletion background, PleC-mCherry exhibited a diffuse localization pattern (Fig. 1C). Expression of sfGFP-PodJΔIDR accumulated at the cell poles in *C. crescentus* (Fig. 1C). However, sfGFP-PodJΔIDR was unable to recruit PleC-mCherry to the cell poles (Fig. 1C). Our observation is consistent with previous PodJ domain analysis in *Caulobacter crescentus*^21^ which indicated a portion of the IDR of PodJ contributes to PleC’s new cell pole localization in *C. crecentus*^7,12^. Therefore, PodJ serves as a scaffold that recruits PleC as a client through its IDR.

### PleC localizes at the cell pole via its tandem PAS sensor

We next asked which domains within PleC serve as the site of interaction with PodJ. PleC contains a periplasmic Cache domain, two cytoplasmic PAS domains in tandem, and a histidine kinase domain. Thus, we heterologously expressed a set of PleC-mCherry domain deletion variants and YFP-PodJ in *E. coli* (Fig. S2B). We observed that PleC-mCherry variants that lack the periplasmic Cache domain or the HK domains co-localized with YFP-PodJ. In contrast, PleCΔPAS-AB-mCherry displayed a diffuse localization pattern. This suggests that PleC’s PAS-AB is required for recruitment to the cell pole by PodJ in *E. coli*.

To corroborate the importance of PAS-AB, we examined the subcellular localization of PleCΔPAS-AB-mCherry or PleC-mCherry in a *podJ* and *pleC* deletion *C. crescentus* background. We observed that the PodJ-PleC complex co-localized at the new cell pole in pre-divisional cells (Fig. 1D). In contrast, in *C. crescentus* strains expressing sfGFP-PodJ and PleCΔPAS-AB-mCherry, the PleC variant displayed a diffuse subcellular localization pattern. This suggests that the PleC-PAS-AB: PodJ-IDR protein-protein interaction is also critical for PleC recruitment to the cell pole in *C. crescentus* (Fig. 1E).

To determine if the PodJ-PleC interaction is conserved across alpha-proteobacteria that encode both PleC and PodJ, we heterologously expressed PodJ and PleC homologs from *Agrobacterium tumefaciens* (Atu), and *Hyphomonas neptunium* (Hyp) individually and together in *E. coli* (Fig. 1F). Both YFP-AtuPodJ and YFP-HypPodJ accumulated at the cell poles in *E. coli*. While heterologous expression of AtuPleC-CFP or HypPleC-CFP individually resulted in a diffuse localization pattern. However, each PodJ homolog co-localized with its cognate PleC at the cell pole (Fig.1F) upon co-expression. Therefore, the ability of PleC and PodJ to co-localize at the cell poles in *E. coli* is conserved amongst these species. Within a representative set of 11 alphaproteobacterial species, we found that the presence of *pleC’s* cytoplasmic PAS-A and PAS-B sensory domains correlates with the existence of a *podJ* and *pilA* homologs in the genome (Fig. S3). This co-conservation agrees with the requirement of PAS-AB for PleC’s recruitment to the cell poles (Fig. 1D, 1F, S2B).

In addition, we found that each of the studied PodJ homologs contains a putative IDR (Fig. S4). These IDRs vary in length from 242 to 261 residues and are flanked by coiled-coil rich regions (Fig. S4A). While the sequences of the IDR are non-conserved, each IDR is composed of two distinct regions (Fig. S4B). One IDR region is rich in negative charges with pI (isoelectric point) between 3.4 and 3.9. In contrast, an adjacent IDR within each PodJ homolog is rich in positive charges with pIs between 9.9 and 10.9. This conservation of charge pattern within the IDRs may play a role in PodJ function.

### Design of a PleC-CcaS chimeric histidine kinase

Due to PodJ interacting with the PleC’s sensory domains (Fig. 1, S2), we hypothesized that PodJ might also regulate PleC function. Complete reconstitution of the PleC signaling network would require the addition of PodJ, PleC, DivK, DivL, CckA, ChpT, CtrA, and a CtrA regulated promoter. We, therefore, leveraged the demonstrated technique of exchanging kinase sensory domains to construct a chimeric histidine kinase to simplify our approach^27,28^. We applied this approach using the green light-sensing CcaS-CcaR system^29^ to interrogate the responsiveness of PleC’s sensor to PodJ expression.

We designed, built, and tested a PleC-CcaS chimeric histidine kinase library (Fig. S5). Each variant consisted of PleC’s tandem PAS sensor fused to different junction sites within the histidine kinase domain of CcaS (Fig. S5A, 2A-2B). *The pleC-ccaS* chimera was co-expressed with its cognate response regulator *ccaR.* Upon phosphorylation, phosphor-CcaR activates transcription of *mCherry* from the *cpcG2* promoter (Fig. 2B). We found that the YFP-PleC-CcaS retained its ability to co-localize with CFP-PodJ in *E. coli* (Fig. 2C). In the absence of PodJ stimulation, the PleC-CcaS chimera exhibited little mCherry expression. In comparison, mCherry expression was highest using chimera AB-1 upon co-expression with PodJ (Fig. S5B). Therefore, the engineered PleC-CcaS chimera retains the on-switch function (Fig. 2D) of the parent green light-sensing CcaS^29^.

**Figure 2:**
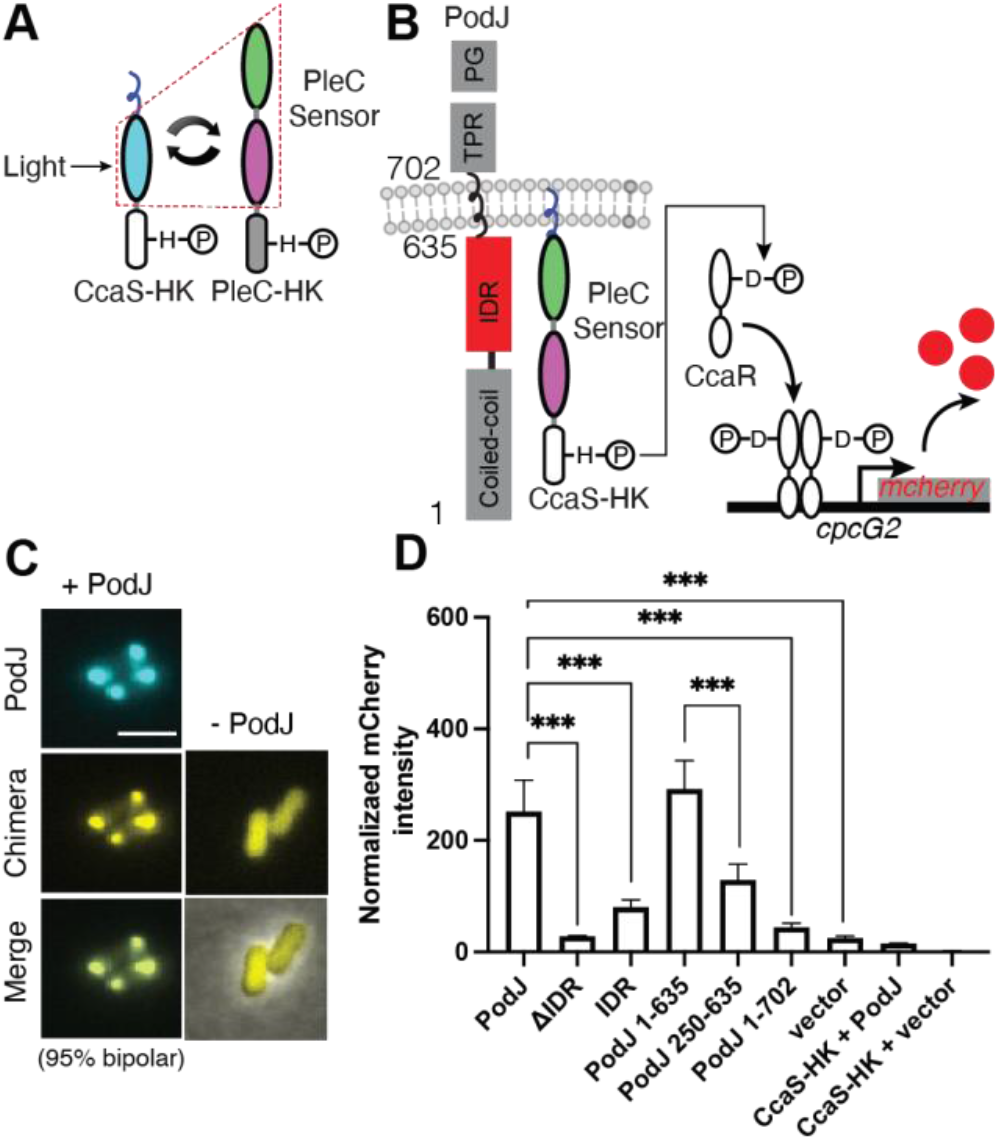
PodJ stimulates the kinase activity of the PodJ-CcaS chimeric histidine kinase in *E. coli*. (A) Design of a PleC-CcaS chimera reporter system. PleC’s cytoplasmic sensory domains were fused to the histidine kinase domain of CcaS. The LOV domain was swapped with PleC’s PAS-AB domain to create a PleC-CcaS chimeric histidine kinase. (B) Upon interaction with PodJ, the PleC-CcaS chimeric histidine kinase phosphorylates CcaR, which then stimulates the expression of mCherry via binding to the *cpcG2* promotor. (C) Heterologous co-expression of CFP-PodJ together with YFP-PleC-CcaS in *E. coli.* CFP-PodJ was induced with 0.5 mM IPTG for 3 hours, and PleC-CcaS-YFP was constitutively expressed. Scale bar: 2 μm. (D) Co-expression of the PleC-CcaS chimera gene reporter system together with PodJ variants. A two-tailed t-test was performed. (n.s: P>0.05, *: P ≤ 0.05, **: P ≤ 0.01, ***: P ≤ 0.001). Error bars represent the standard deviation from three independent biological replicates performed on different days.

The PleC-CcaS chimera AB-1 connects PAS-AB to CcaS between the D and V of the conserved DVT hinge motif^30^ (Fig. S6A-C). These results suggest that we have engineered a functional PodJ-responsive PleC-CcaS chimeric histidine kinase.

### Stimulation of PleC-CcaS activity requires PodJ’s IDR

To determine whether PleC-CcaS stimulation depended upon specific interactions with PodJ, we compared the effects of co-expressing the YFP-PodJ domain deletion variants with the PleC-CcaS chimera (Fig. 2D). Before the analysis, we confirmed that the YFP-PodJ variant displayed fluorescence over 10-fold greater than the empty vector (Fig. S7A). Relative to the empty vector control, the expression of PodJ stimulated the PleC-CcaS chimera and resulted in a 10-fold increase in mCherry expression (Fig.2D). However, a PodJ variant that lacks the IDR, YFP-PodJ-ΔIDR, was unable to stimulate PleC-CcaS-mediated mCherry expression. In contrast, expression of the YFP-IDR alone led to a 3-fold activation of mCherry expression (Fig.2D). This suggests that stimulation of the PleC-CcaS chimera is dependent upon the interaction with PodJ’s IDR. However, full stimulation of PleC-CcaS requires the entire cytoplasmic domain, suggesting a role of the coiled-coils in PleC-CcaS regulation.

### PodJ_S_ is unable to stimulate the PleC-CcaS chimera

Next, we asked if the PleC-CcaS chimera could be stimulated by PodJ_S_, the primary form in *C. crescentus* swarmer cells. Past experiments indicated that loss of the periplasmic domain leads to reduced expression of CtrA mediated genes in *C. crescentus*^7^. We observed that PodJ_L_ and PodJ_S_ could recruit PleC to the cell poles when heterologously expressed in *E. coli* (Fig. S2A), consistent with the past observation in *C. crescentus*^6^. Therefore, the down-regulation of the CtrA pathway upon PodJ proteolysis is not due to a loss of PleC-PodJ binding. Unlike the expression of full-length PodJ_L_ we observed that expression of PodJ_S_ was unable to stimulate mCherry expression via the PleC-CcaS chimera (Fig. 2D).

The loss of PodJ_S_ stimulation of PleC could result from an altered PodJ transmembrane anchoring that allosterically affects the IDR-PAS conformational state. Therefore, we also examined if PodJ variants lacking the TM tether (PodJ(1-635)) could relieve steric effects on the IDR-PAS conformation. We observed that the cytoplasmic PodJ variant lacking the TM tether stimulated mCherry expression in the PleC-CcaS reporter assay (Fig. 2D). This indicates that PodJ’s cytoplasmic domains alone can stimulate PleC-CcaS function. Meanwhile, it suggests that the transmembrane region may regulate the PodJ-PleC signaling complex, consistent with the reduced CtrA pathway activation of strains expressing PodJ_S_ in *C. crescentus*^7,21^.

We next asked how cell pole accumulation of the PodJ-PleC complex impacted PleC-CcaS signaling. Construct lacking the N-terminal coiled coils, PodJ(250-635), does not accumulate at the cell poles in *E. coli* (Fig S2) or *C. crescentus* (Fig. S1). We found that PodJ(250-635) led to a 3-fold increase in mCherry expression compared to PodJ(1-635), which stimulated a 10-fold increase. These results indicate cell pole accumulation of PodJ is not required for PleC-CcaS stimulation but may impact the degree of PleC-CcaS stimulation.

### Stimulation of PleC-CcaS requires functional PAS domains

Given that PodJ recruits PleC through its cytoplasmic tandem PAS sensory domains, we hypothesized that the regulation of PleC-CcaS kinase domains requires sensory domain stimulation. Notably, our sequence analysis indicated that PleC’s PAS A and B domains share low similarities (47.88%) (Fig. S6A). In contrast, PAS-A homologs across alphaproteobacteria exhibit 68.3% sequence similarity, while PAS-B homologs exhibit 64.4% (Fig. S6D). Therefore, we suspect that each PAS domain may have a distinct function.

Previous studies have shown that the signal transmission motif D-I/V-T at the C terminus of the PAS domain allosterically relays signals from the central binding cavity to the C-terminal coiled-coil linker^31^. These studies showed that mutations of the Asp to Ala or the Thr to Ala within this motif disrupted the PAS sensor’s sensitivity to signal stimulation^28,32^. Similarly, we hypothesized that PodJ’s stimulation of PleC activity also requires PleC’s PAS-A and PAS-B conserved motifs^31^ (Fig. S6).

We, therefore, generated PAS-A (D433A, V435A) and PAS-B (D548A, V550A) hinge motif PleC-CcaS variants to disrupt signal flow within each PAS domain (Fig.3A). Before analysis, we confirmed the expression of PodJ by measuring the fluorescence intensity of N-terminal YFP (Fig. S7B). The wild-type PleC-CcaS chimera displayed a 10-fold increase in mCherry expression upon PodJ induction. In contrast, PleC-CcaS PAS-A and PAS-B variants showed little to no stimulation by the expression of PodJ_L_ (Fig. 3B). Also, compared to wild-type, the changes to the PAS domain motif appear to lock the downstream kinase into a state with a low activity which was insensitive to PodJ expression. Thus, the reduction of stimulation by PodJ could be due to a loss of PAS domain allostery or loss of PodJ-PleC co-localization. Moreover, each PAS domain should be functional to mediate PodJ stimulation of PleC-CcaS.

**Figure 3:**
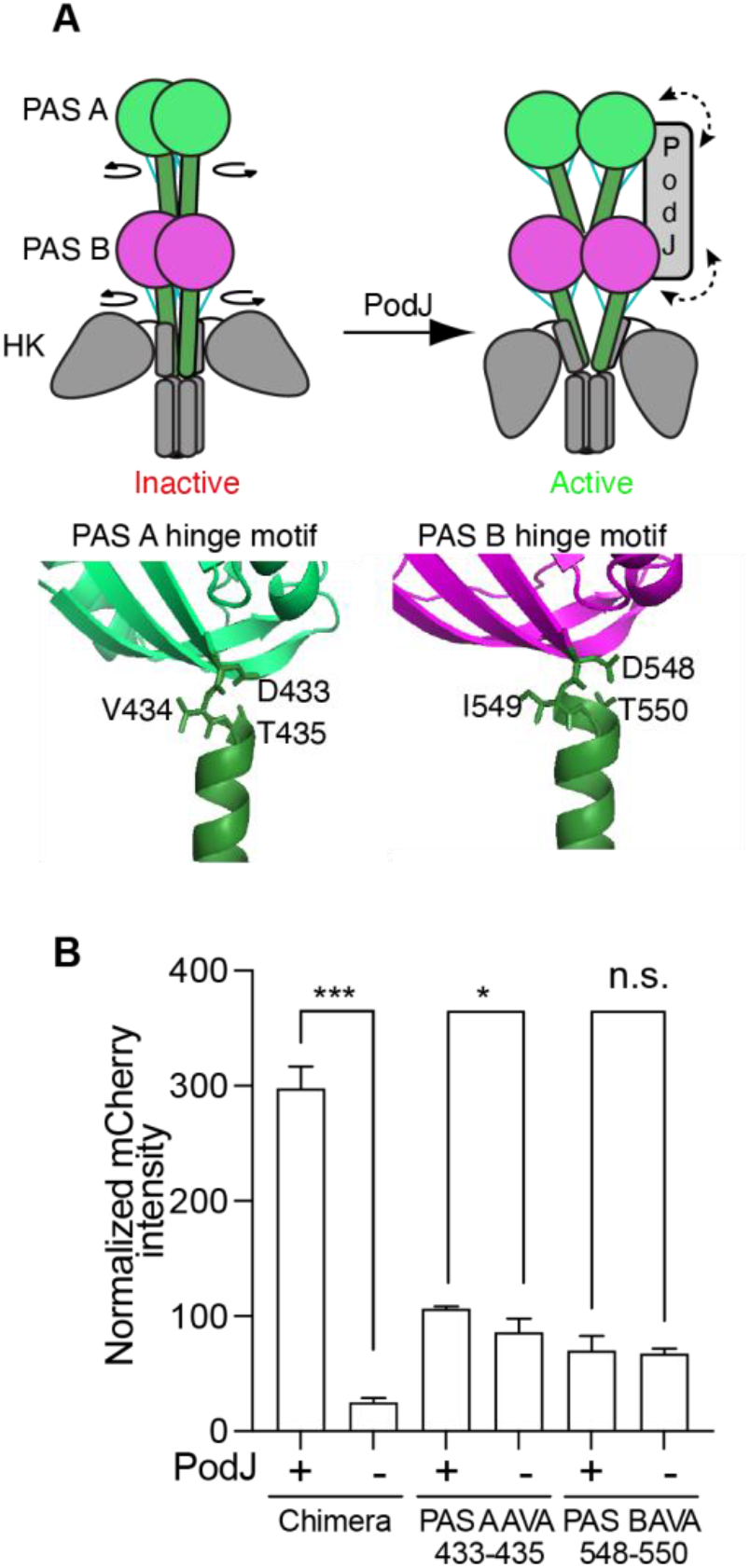
Stimulation of the PleC-CcaS chimera by PodJ depends on the signaling transmission motifs at the C-terminus of PleC PAS-A and PAS-B. (A) Cartoon of the conformational change that occurs at the signal transmission motif of sensor kinases. The signal transmission motifs between PAS-A (green), PAS-B (magenta), and the coiled-coil linker (green) contain residues that form several hydrogen bonds (blue) and serve as a conformational switch. Homology model of PleC compared to YF1 (PDB ID: 4GCZ-A)^32^.In the left cartoon, a curled arrow around the end of each PAS domain indicates the relative rotation of the N-terminal linker upon signal stimulation. The bidirectional dashed arrow in the right cartoon suggests the interaction between PodJ and PleC PAS sensory domains. (B) Mutation of the PAS-A (D132A, T134A) or PAS-B (D247A, T249A) PAS sensor transmission motif results in PleC-CcaS mutant chimeras that are unresponsive to PodJL. Error bars represent the result of three independent biological replicates. A two-tailed t-test was performed. (n.s: P>0.05, *: P ≤ 0.05, **: P ≤ 0.01, ***: P ≤ 0.001)

### PodJ forms biomolecular condensates *in vitro*

To reconstitute the PodJ-PleC signaling system *in vitro,* we utilized PodJ(1-635) as it accumulated the cell poles (Fig. 1B, S2A) and stimulated PleC-CcaS signaling (Fig. 2). By size-exclusion chromatography and native gel analysis, we observed that PodJ oligomerized as a high molecular weight oligomer (>670 kDa) (Fig. 4A, 4B). This large oligomer size could be poised to mediate weak multivalent interactions that promote phase separation. Indeed, we observed *in vitro* that the sfGFP-PodJ(1-635) protein phase separates as round protein-rich droplets with a saturation concentration between 1.5 and 2.0 μM (Fig. 4C, S8A). This C_sat_ is less than the estimated PodJ concentration in cells from ribosome profiling measurements of 2-5 μM^33^. Moreover, time-lapse imaging revealed that the PodJ droplets fuse upon contact and gradually relax back to a spherical droplet over 12 minutes, demonstrating liquid-like properties *in vitro* (Fig. 4D).

**Figure 4.**
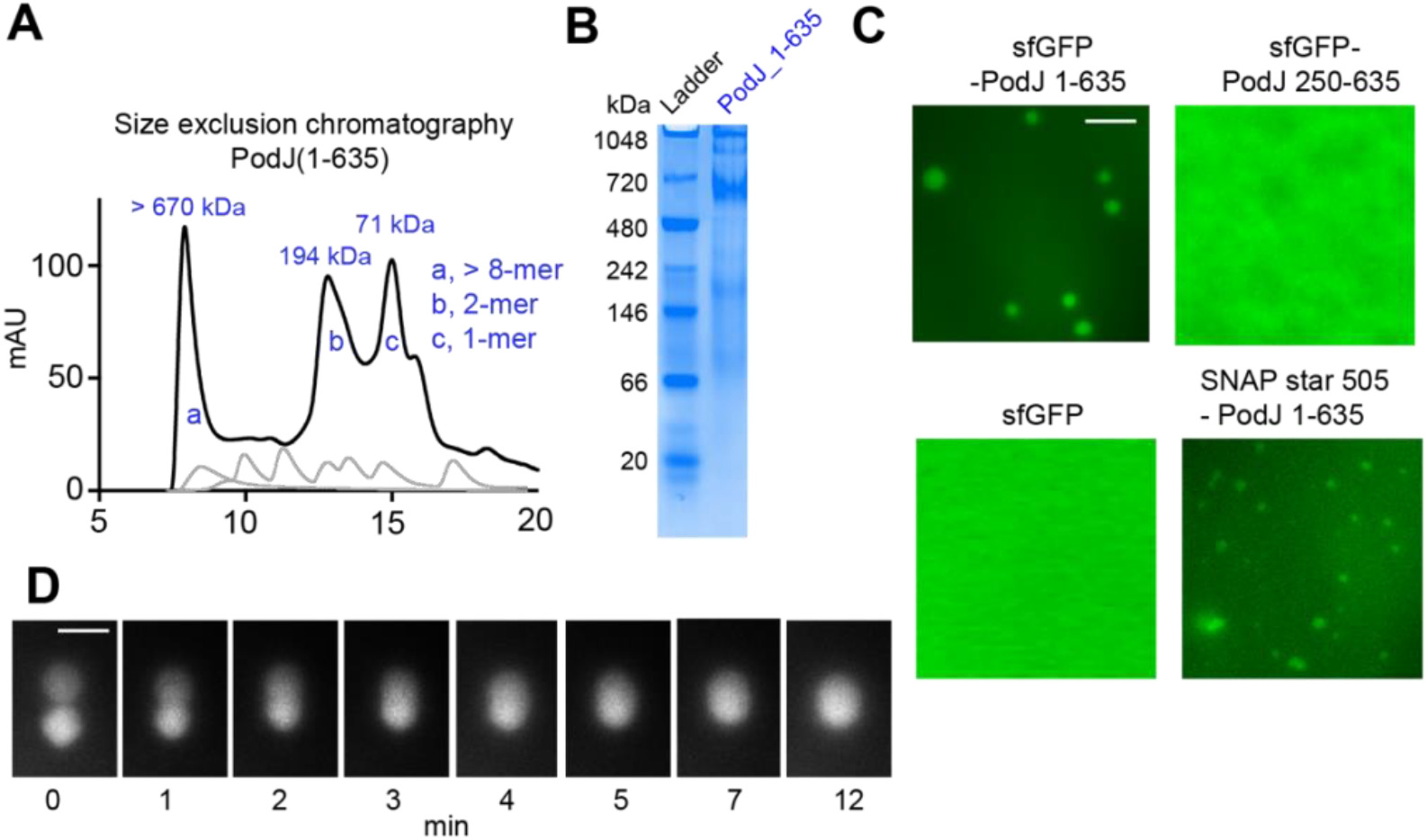
PodJ phase separates as a biomolecular condensate *in vitro.* The oligomerization state of PodJ(1-635) was analyzed via (A) size exclusion chromatography and (B) native gel. (C) Fluorescence microscopy images of purified sfGFP-PodJ(1-635), sfGFP-PodJ(250-635), sfGFP and SNAP-PodJ(1-635) at 7.5 μM, 50 mM Tris-HCl pH 8.0 and 200 mM KCl. Concentration-dependent assembly of each construct is reported in Fig. S8. (D) Time-lapse imaging of individual PodJ droplets undergoing dynamic liquid-like fusion events. Scale bar: 2.5 μm.

To determine the impact of fluorescent protein interactions, we observed that sfGFP did not form any visible droplets (Fig. 4C, S8B). In addition, SNAP-tag fused to the N-terminus PodJ(1-635) also formed droplets at the same condition with a smaller size (Fig. S8C). These results indicate that PodJ’s droplet formation does not require sfGFP. However, weak fluorescent protein interactions likely mediate increases in protein droplet size.

To interrogate the biomolecular condensate material properties of PodJ, we examined the impact of 1,6-hexanediol (HD), which is commonly used to disrupt weak hydrophobic interactions and dissolve biomolecular condensates^34^. We observed that the addition of 10% HD disrupted droplet formation (Fig. S9A). Moreover, adenosine triphosphate (ATP) has been shown to act as a biological hydrotrope that dissolves liquid-liquid phase separation (LLPS) assemblies^35, 9^. Similarly, we observed that the addition of 10-20 mM ATP or ADP led to a reduction in droplet size, partitioning ratio and PleC-mCherry recruitment (Fig. S9B-C). In contrast, we observed that the addition of physiological concentrations of 125-1000 μM ATP led to increased sfGFP-PodJ droplet size and partitioning (Fig. S9B, D). This increase in droplet size and partitioning may suggest ATP binding to PleC and subsequent biochemical activities may influence droplet size. In summary, the PodJ biomolecular condensate assembly properties are tunable by adding common small-molecule LLPS regulators.

We next considered if the ability to localize at the cell poles correlated with PodJ’s ability to phase separate *in vitro*. Therefore, we purified and analyzed sfGFP-PodJ(250-635), which lacks CC1-3 (Fig. 4C). We observed sfGFP-PodJ(250-635) increased the critical assembly concentration to between 3.5 to 4.0 μM (Fig. 4C, S8A), and about 2-fold higher than sfGFP-PodJ(1-635). Therefore, the N-terminal coiled-coil domains are needed for robust PodJ phase separation *in vitro* (Fig. 4C, S8A) and mediate PodJ cell pole accumulation *in vivo* (Fig. 1A).

### PodJ biomolecular condensate recruits PleC *in vitro*

We then considered if the *in vitro* sfGFP-PodJ(1-635) biomolecular condensates could recruit PleC-mCherry as a client. We observed that PleC-mCherry could readily accumulate within the PodJ-rich droplets, while mCherry alone was not enriched (Fig. 5A). In contrast, mCherry, mCherry + sfGFP, PleC PAS AB-HK-mCherry, and mCherry + sfGFP-PodJ 1-635 did not form droplets under the same condition (Fig. S8B).

**Figure 5.**
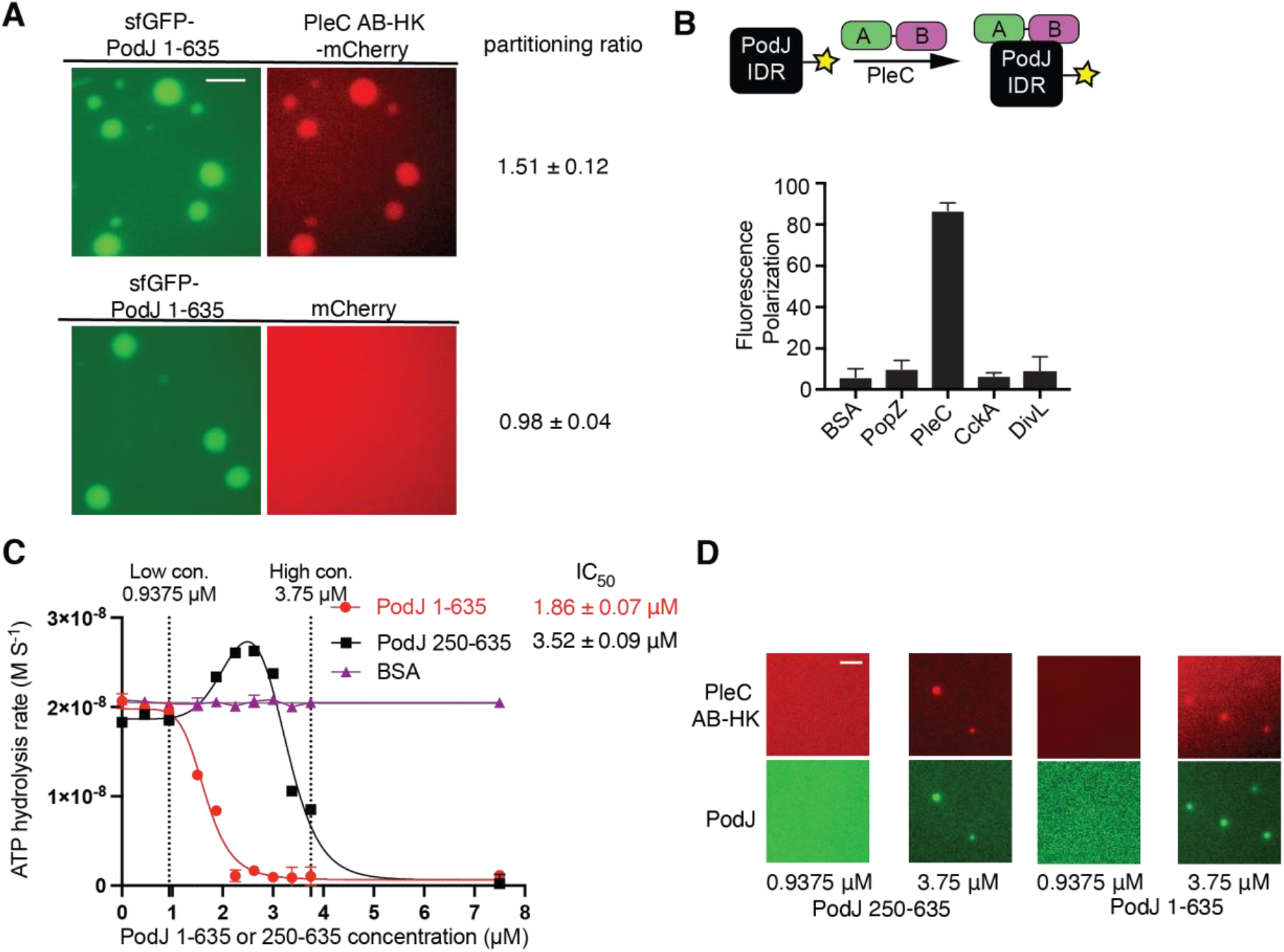
In vitro PodJ recruits PleC as a client and represses its kinase activity. (A) Fluorescence microscopy images of purified sfGFP-PodJ with PleC-mCherry. Purified proteins were mixed at 7.5 μM, 50 mM Tris-HCl pH 8.0 and 200 mM KCl. Scale bar: 2.5 μm. (B) Fluorescence polarization binding assay of BODIPY dye-labeled 250 nM PodJ-IDR mixed with the following: 10 μM BSA, PopZ, PleC PAS-AB, CckA (70-691), or DivL (54-769). (C) Coupled enzyme assay measures the PodJ-activated activity switch of PleC. Conditions from left to right: 7.5 μM of PleC PAS AB-HK was incubated with 1 mM ATP and 0, 0.45, 0.9375, 1.875, 2.25, 2.625, 3, 3.375, 3.75, 7.5 μM of PodJ(1-635) (red circle), PodJ(250-635) (black square) or BSA (shown in purple square), respectively. Data was fitted through non-linear regression into [inhibitor] versus response for PodJ(1-635) and bell-shaped for PodJ 250-635 using Prism 9. Error bars represent the standard deviation from three independent biological replicates. (D) Fluorescence microscopy images of kinase reaction mixtures containing purified sfGFP-PodJ(1-635) and sfGFP-PodJ(250-635) at 0.9375 μM and 3.75 μM. Purified proteins were mixed in kinase buffer supplemented with 1.0 mM ATP, 10 mM MgCl_2_, 3 mM phosphoenolpyruvate, 0.2 mM NADH, 2 units of pyruvate kinase, and 6.6 units lactate dehydrogenase. Scale bar: 2.5 μm.

We employed an *in vitro* fluorescence polarization assay to detect the interaction between PodJ’s IDR and histidine kinases co-localized with PodJ at the new cell pole, including PleC PAS-AB domains, DivL, and CckA. We fluorescently labeled PodJ’s IDR with a BODIPY dye and observed robust binding between PleC and PodJ’s IDR. In contrast, we could not detect any interactions between PodJ’s IDR and DivL and CckA (Fig. 5B). These fluorescence polarization studies indicate that PleC’s tandem PAS domains interact specifically with the IDR from PodJ.

### PodJ represses PleC kinase activity *in vitro*

To evaluate the impact of PodJ(1-635) upon PleC, we employed a coupled enzyme assay to detect changes in kinase activity^36^. PleC PAS AB-HK at 7.5 μM supplemented with 1000 μM ATP displayed a 2 × 10^−8^ M s^−1^ ADP production rate (Fig. 5C). We observed dose-dependent repression of the ADP production rate upon addition of PodJ(1-635) to PleC PAS AB-HK. We also found that the addition of 2.5 μM PodJ(1-635) with to PleC PAS AB-HK at 7.5 μM supplemented with 1000 μM ATP displayed a 1.1 × 10^−9^ M s^−1^ ADP production rate with an IC_50_ of 1.86 ± 0.07 μM.

We next asked if the diminished phase separation capabilities of PodJ(250-635) impact its catalytic capabilities. We observed that the ADP production IC_50_ for PodJ(250-635) was two-fold higher than PodJ(1-635). Interestingly, our imaging of sfGFP-PodJ(1-635) or sfGFP(250-635) indicated that low concentrations of PodJ protein do not phase separation and do not repress PleC kinase activity. In contrast, higher PodJ concentrations of both constructs lead to phase separation and repression of PleC kinase activity (Fig. 5D). However, the addition of BSA resulted in no change in PleC ATP hydrolysis activity (Fig. 5C). Therefore, the interaction of PodJ with PleC specifically represses PleC’s histidine kinase function. While kinase activity repression typically correlates with increased phosphatase function^10,37,38^, future biochemical studies should examine how PodJ impacts PleC’s ATP binding, phosphatase and phosphotransfer functions.

Notably, *in vivo*, PleC-CcaS chimera is inactive when expressed alone, and the addition of PodJ(1-635) stimulated kinase activity (Fig. 2D). By contrast, *in vitro*, PleC exhibited kinase activity in solution alone, but the addition of PodJ(1-635) represses PleC kinase activity (Fig. 5C). The differences in regulation are likely rooted in two aspects of the PleC-CcaS design. First, truncation of PleC’s periplasmic sensor may impact the structure of the cytoplasmic sensor. Past studies have shown that alterations of the N or C-terminus of PAS sensory domains can impact sensor functions as an activator or repressor upon stimulation^32,39^. Similarly, we suspect that N or C-terminal modifications to PleC’s sensor have switched its mode of regulation compared to the wild-type PleC HK (ON versus OFF-switch). Second, Ple-CcaS chimeric histidine kinase adopts the same on-switch behavior as the green light-sensing CcaS histidine kinase. Indeed, the histidine kinase domains of CcaS are divergent from PleC’s histidine kinase domain and may utilize different structural mechanisms. Nevertheless, both experimental sets provide evidence that PleC’s tandem sensory domain serves as a sensor for PodJ.

## Discussion

Reconstitution of the PodJ-PleC complex *in vitro* (Fig. 4-5) and in *E. coli* (Fig. 1-3) demonstrated that PodJ recruits and regulates PleC function through a PAS-IDR interaction. Our studies demonstrated that *in vitro* PodJ-PleC phase separates as a biomolecular condensate. Therefore, we propose that the PodJ forms a biomolecular condensate that compartmentalizes and regulates PleC at the cell poles. We showed that the N-terminal coiled-coils were critical for cell pole accumulation of PodJ’s cytoplasmic domains (Fig. 1B). Future studies will consider how PodJ’s membrane tethering and its periplasmic domains impact PodJ phase separation. Moreover, additional studies are needed to conclude if PodJ’s ability to phase separate *in vivo* influences its ability to accumulate at the cell poles.

The discovery of PodJ’s phase separation capabilities and ability to directly regulate PleC function provides new data to understand how PodJ is logically connected to the cell cycle. Past work demonstrated that the cell-cycle master regulators control PodJ’s expression^40^ and proteolysis once per cell-cycle^22,23^. We observed that the shortened form of the PodJ scaffold retains the ability to phase separate and recruit PleC to the cell poles. However, PodJ_S_ no longer stimulates PleC-CcaS function. Thus, proteolytic remodeling of the PodJ within these biomolecular condensates serves as a cell-cycle checkpoint signal to tune PleC function (Fig. 6).

**Figure 6.**
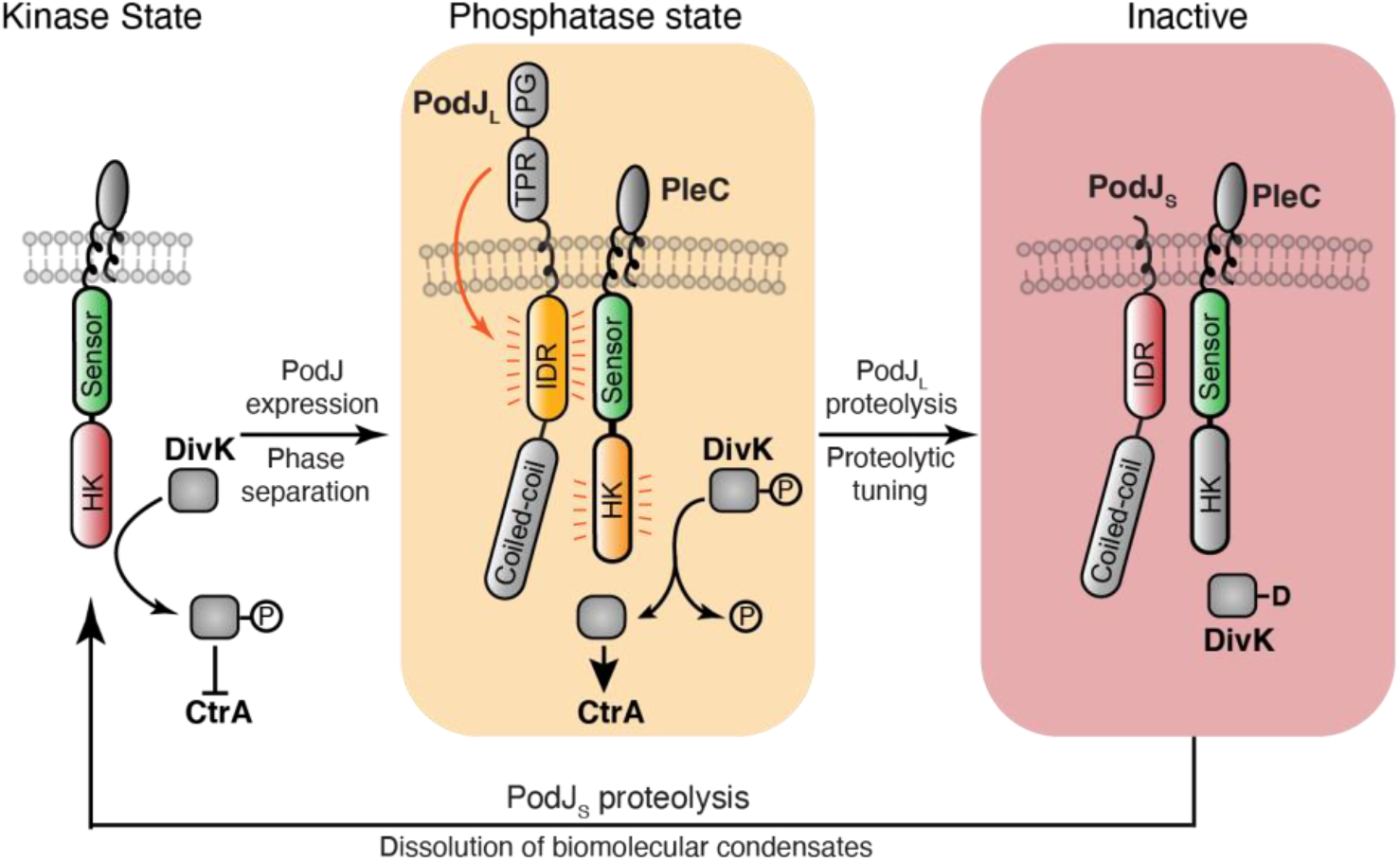
Proposed model for cell-cycle dependent phase separation, proteolytic tuning of biochemical function, and dissolution of a biomolecular condensate that regulates a histidine kinase. PodJ serves as a cell-cycle checkpoint signal that regulates the formation and activity of PodJ-PleC biomolecular condensate. PleC alone can function as a kinase, which leads to inhibition of the CtrA pathway and expression of PodJ_L_. The PodJ_L_-PleC complex phase separates and represses PleC kinase activity. This leads to CtrA pathway activation and the expression of PodJ proteases. Proteolysis of PodJL to its short form PodJ_S_ leads to an inactive form of PleC that may have kinase functions. Proteolysis of PodJ_S_ leads to the dissolution of the PodJ-PleC biomolecular condensate and liberates PleC from the cell pole.

Upon complete degradation of PodJ_S_, this biomolecular condensate is dissolved and PleC is liberated from the cell pole. In this diffuse state, PleC’s activity is regulated by DivK allosteric stimulation^17^ and pilus retraction^18^. PleC kinase function stimulates down-regulation of the CtrA pathway and the expression of new PodJ protein at the swarmer-to-stalk transition^40^. This leads to stimulation of new cell pole localized PleC phosphatase function and activation of the CtrA pathway. Subsequently, robust activation of the downstream CtrA signaling pathway leads to the expression of PodJ specific proteases^22,23^ that proteolyze PodJi_ into PodJ_S_. This proteolytic event tunes the functions of the PodJ biomolecular condensates resulting in phosphatase downregulation.

More broadly, phase separation provides an accessible compartmentalization strategy for bacterial kingdom^41^. The three scaffolding proteins, PopZ^8^, SpmX^9^, and PodJ (Fig. 4-5), phase separate as two distinct membraneless organelles at opposite ends of the cell. In addition, several other recent studies have shown that bacteria leverage phase separation to compartmentalize and regulate RNA polymerase^42^, mRNA decay^43,44^, ABC transporters^45^, DNA repair^46^, cell division^47^, chromosome segregation^48^ and carboxysome assembly^49^. These studies have laid the foundation to consider the extent and advantages of organizing biochemistry within biomolecular condensates in the bacterial cytoplasm.

Biomolecular condensates are generally thought to enhance enzyme reaction rates through mass action^50^. Some recent studies have shown enhanced enzymatic rates, including RNA decapping^51^, pyrenoid biochemistry^52^ and SUMOylation^53^. Here we have examined how biomolecular condensates regulate the histidine kinase activity of PleC. To determine the role of allostery, we found that a functional PAS sensor transmission motif was required for stimulation by PodJ (Fig. 3). This indicates that PodJ stimulation requires PleC-CcaS’s sensor to kinase domain allosteric stimulation. Interestingly, a comparison of PodJ-IDR vs. full-length PodJ stimulation of PleC-CcaS revealed a 3-fold versus 10-fold degree of stimulation. This additional 3.3-fold activation suggests a unique additional contribution from PodJ’s phase-separated chemical environment and mass action.

Similarly, we observed that PodJ could repress the activity of PleC *in vitro* under conditions where phase separation readily occurred. In both cases, we suspect that there are interconnected contributions from mass action and kinase allostery. The localized high concentration of PodJ-PleC may promote an active PodJ-PleC conformation in contrast to the lower PodJ-PleC concentration in the dilute phase. Similarly, it was shown that phase separation of the *Saccharomyces cerevisiae* RNA decapping complex Dcp1/Dcp2 was coupled to conformational control of RNA decapping enzymatic activity^51^. Two other histidine kinases, CckA^10^ and DivJ^9^, also show activation of enzyme function at high packing density on liposomes. In addition, both PleC and CckA require their N-terminal PAS sensory domain to mediate enzyme regulation at increased local concentration. Collectively these three studies suggest strategies that concentrate histidine kinases, such as phase separation, may have a generalizable effect of robustly stimulating low copy histidine kinases^9,10^.

In summary, two-component systems provide bacterial cells with a customizable signaling platform. PleC has been customized for spatial control through phase separation and temporal control through sensing the cell-cycle produced and degraded signal PodJ. Moreover, PleC integrates both intra- and extracellular signals^18^ to coordinate growth and development. Therefore, the interplay of scaffolds, phase separation and two-component systems provide avenues to orchestrate complex development in bacteria.

## Methods

### Bacterial Strains

All experiments were performed using *Caulobacter crescentus* NA1000 (also known as CB15N) and *Escherichia coli* DH5α and BL21 purchased from Promega. *C. crescentus* NA1000 was a kind gift from Dr. Lucy Shapiro (Stanford University School of Medicine). PCR reactions and primers used for Gibson assembly are listed in Table S1. All relevant primers are given in detail in Table S2. Plasmids used in this study are listed in Table S3. More strains are listed in Table S4. Transformations were carried out as described^54^. Details about plasmid and strain construction are listed in Text S1.

### Plasmid cloning strategies

Fragments of target genes plasmid backbone were amplified via PCR using Phusion polymerase (Thermo Scientific). PCR reactions were performed in 50 μL reaction mixtures containing 3% (v/v) DMSO, 1.3 M betaine, 0.3 μM each primer, 0.2 mM each dNTP, and 1U Phusion High-Fidelity DNA Polymerase (Thermo Scientific). Both fragments were purified via gel extraction. Gibson assembly^55^ reactions were performed in 20 μL with 100 ng backbone and typically a 1:10 backbone: insert ratio. A Gibson reaction master mix was prepared from 5x reaction buffer, T5 exonuclease (NEB), Phusion polymerase (NEB), Taq ligase (NEB), and stored as aliquots of 15 μl at −20°C. An annealing temperature of 55°C was used for all reactions, followed by 10 min at 4°C and 10 μl of the reaction product was then transformed into chemically competent *E. coli* DH5a cells using the KCM transformation method. Oligonucleotide primers applied for amplification of the gene insert were designed using the j5 online program, and they featured overlaps of 26 bases to the insertion site in the plasmid.^56^. Oligonucleotides were synthesized by IDT (Coralville, IA), and all DNA sequencing reactions were performed by Genewiz (South Plainfield, NJ). DNA oligos, plasmid construction methods, plasmids, and strains used in this study are listed in Table S1-S4.

### Growth and Induction Conditions

*C. crescentus* strains were grown at 28°C in PYE (peptone yeast extract). *E. coli* strains used for protein purifications and microscopy experiments were grown at 37 °C in LB medium unless otherwise stated. When required, protein expression was induced by adding 0.002-0.5 mM Isopropyl β-D-1-thiogalactopyranoside (IPTG) or 0.5-10 mM arabinose in *E. coli* 0.003%-0.3% xylose or 0.005-0.5 mM vanillic acid in *C. crescentus* unless otherwise stated. The induction time for microscopy experiments is 0.5-2 hours in *E. coli* and 3-5 hours in *C. crescentus*.

### Phase Contrast and Epifluorescence Microscopy

Cells were imaged after being immobilized on a 1.5% w/v agarose/media (PYE for *C. crescentus* and LB for *E. coli*) pad. Phase microscopy was performed by a Nikon Eclipse T*i*-E inverted microscope equipped with an Andor Ixon Ultra DU897 EMCCD camera and a Nikon CFI Plan-Apochromat 100X/1.45 Oil objective. The excitation source was a Lumencor SpectraX light engine. Chroma filter cube CFP/YFP/MCHRY MTD TI was used to image ECFP (465/25M), EYFP (545/30M), and mCherry (630/60M). Chroma filter cube DAPI/GFP/TRITC was used to image sfGFP (515/30M). Images were collected and processed with Nikon NIS-Elements AR software, ImageJ^57^, and MicrobeJ^58^.

### Protein expression and purification of PodJ, PopZ, PleC, CckA, and DivL

Protein expression for PodJ, PopZ, PleC, and CckA followed the same protocol described in detail below for PodJ(1-635). To purify the cytoplasmic portion of PodJ(1-635), Rosetta (DE3) containing plasmid pwz091 was grown in 6 liters LB medium (30 μg/ml chloramphenicol and 100 μg/ml ampicillin) at 37°C. The culture was then induced at an OD_600_ of 0.4–0.6 with 0.5 mM IPTG overnight at 18°C. The cells were harvested, resuspended in the lysis buffer (50 mM Tris-HCl, 700 mM KCl, 20 mM Imidazole, 0.05% dextran sulfate, pH 8.0) in the presence of protease inhibitor cocktail tablets without EDTA (Roche). The cell suspension was lysed with three passes through an EmulsiFlex-C5 cell disruptor (AVESTIN, Inc., Ottawa, Canada), and the supernatant was collected by centrifuging at 12 000 *g* for 30 min at 4°C. In addition, the insoluble cell debris was resuspended by the recovery buffer (50 mM Tris-HCl, 1000 mM KCl, 20 mM Imidazole, 0.05% dextran sulfate, pH 8.0), and its supernatant was collected as well as the previous centrifugation. The combined supernatants were loaded onto a 5 ml HisTrap^™^ HP column (GE Healthcare) and purified with the ÄKTA^™^ FPLC System. After washing with 10 volumes of wash buffer (50 mM Tris-HCl, 300 mM KCl, and 25 mM imidazole, pH 8.0), the protein was collected by elution from the system with elution buffer (50 mM Tris-HCl, 300 mM KCl, and 500 mM imidazole, pH 8.0). Then the protein was concentrated to a 3 ml volume using Amicon Centrifugal Filter Units, resulting in over 95% purity. Then the protein was dialyzed with a buffer containing 50 mM Tris-HCl (pH 8.0), 300 mM KCl, and then aliquoted to a small volume (100 μl) and kept frozen at -80°C until use.

### Labeling of SNAP-PodJ(1-635) with SNAP-Cell^®^ 505-Star

SNAP-PodJ(1-635) was purified by following the protocol described above until the FPLC purification and Amicon centrifuge concentration step. Then the protein in the wash buffer was incubated with SNAP-Cell^®^ 505-Star in a 1:1.2 molar ratio at 0 °C for 3 hours. Next, the mixture was dialyzed against a dialysis buffer containing 50 mM Tris-HCl (pH 8.0), 300 mM KCl, then aliquoted to a small volume (20 μl) and kept frozen at -80°C until ready to use for imaging.

### Fluorescence Polarization Assay

To label PodJ-IDR (471-635), we cloned a cysteine just after the 6X-His-tag proteins at the N-terminal of each protein. Cys-PodJ-IDR expression and purification followed the same protocol as PodJ mentioned above. These two proteins were labeled at the cysteine using thiol-reactive BODIPY^™^ FL N-(2-Aminoethyl) Maleimide (Thermo Fisher). The proteins were mixed with 10-fold excess BODIPY^™^ FL N-(2-Aminoethyl) Maleimide and allowed to react for 2 hours at room temperature. The unreacted dye was quenched with mercaptoethanol (5% final concentration). The labeled proteins were purified via dialysis to remove unreacted fluorescent dye (5 times, 500 ml buffer, and 30 min each).

Fluorescence polarization binding assays were performed by mixing 100 nM labeled proteins with 0, 0.25, 0.5, 1, 2, 4, 8, 16 μM partner protein (PopZ, CckA, PleC, DivL, or BSA) for 45 minutes to reach binding equilibrium at the room temperature. Fluorescent proteins were excited at 470 nm, and emission polarization was measured at 530 nm in a UV-vis Evol 600 spectrophotometer (Thermo Fisher). Fluorescent polarization measurements were performed in triplicates, and three independent trials were averaged with error bars representing the standard deviation.

### Quantification and Statistical Analysis

FIJI/ImageJ^57^ and MicrobeJ^58^ were used for image analysis. More than 100 representative droplets were selected for partitioning ratio calculation, and each droplet’s fluorescent intensity inside was divided by the background intensity outside. The mean and standard deviation for each measurement is shown. The number of replicates and the number of cells analyzed per replicate is specified in corresponding legends. All experiments were replicated three times, and statistical comparisons were carried out using GraphPad Prism with two-tailed Student’s t-tests. Differences were significant when *p* values were below 0.05. In all figures, measurements are shown as mean ± standard deviations (s.d.).

### Fluorescence microscopy imaging of Biomolecular Condensates

sfGFP-PodJ(1-635), PleC PAS AB-HK-mCherry, SNAP-PodJ(1-635), sfGFP, mCherry protein aliquots were thawed on ice along with KCl, Tris-HCl (pH=8.0), 1,6-hexanediol, ATP, ADP and sterile water. Working solutions of protein, Tris-HCl (pH=8.0), and KCl were combined, diluted with water to various concentrations, and incubated at room temperature for 15 minutes before imaging. Then the incubated sample mix was pipetted onto the slides with SureSeal Imaging Spacers (Electron Microscopy Sciences) and covered with coverslips (VWR). All images were taken with an Eclipse Ti-E inverted microscope (Nikon) in both phase-contrast and fluorescent channels using a Plan Apo 100x objective.

### Size exclusion chromatography and native gel analysis

A gel filtration standard (Sigma) containing thyroglobulin (bovine, 669 kDa), carbonic anhydrase (bovine, 29 kDa), blue dextran (2,000 kDa), apoferritin (horse, 443 kDa), β-Amylase (sweet potato, 200 kDa), alcohol dehydrogenase (yeast, 150 kDa), and albumin (bovine, 66 kDa) were used to generate a molecular weight standard plot using a Superdex 200 10/300 GL column (GE Healthcare). A 3.2 mg/ml sample of His-PodJ(1-635) was loaded onto the column and eluted after 7.9 ml, 12.8 ml, and 15.0 ml of buffer, corresponding to a molecular weight of 1,851 kDa, 194 kDa, and 70.7 kDa (theoretical monomer = 73.0 kDa). One representative result of triplicates was shown.

His-PodJ(1-635) was also analyzed by running a native gel. Protein was separated by gel electrophoresis (8% resolving gel) at 80 V for 4 hours at 4^°^C, using a native protein ladder (range from 66 to 669 kDa, Thermo Fisher).

### Western blotting

We analyzed protein levels and potential proteolysis for each protein construct expression in *E. coli* and *C. crescentus* through western blot analysis. These assays indicated that each PodJ and PleC variant was expressed and exhibited little to no proteolysis (Fig. S10). For the western blotting, log-phase cells were induced 0.002-0.5 mM Isopropyl β-D-1-thiogalactopyranoside (IPTG) or 0.5-10 mM arabinose in *E. coli* for 0.5-2 hours, and 0.003%-0.3% xylose or 0.005-0.5 mM vanillic acid in *C. crescentus* for 3-5 hours unless otherwise stated. After induction was complete, the cells were pelleted and resuspended in 250ml of 2x Laemmli buffer for each 1.0 OD_600_ unit. The samples were boiled at 95°C for 10 min then vortexed. Next, 10μl of samples were loaded in a 10% SDS-PAGE gel and run at 125V for 90 min. Then the transfer was done at 20V for 80 minutes at 0 °C. Blocking was done for 1 hour using 25ml of blocking buffer (25ml 1x TBST, 1.25g non-fat dry milk) at 0 °C with gentle shaking. For primary antibody blotting, the membrane was submerged in 1:5000 dilution of the anti-GFP (#2956S, Cell Signaling) or anti-mCherry (#43590S, Cell Signaling) antibody in the buffer (10 ml 1x TBST, 0.5g BSA, 10 μl antibody) and shacked gently for 1 hour at room temperature. After washing the membrane 3 times with 1x TBST for 5 minutes each, the membrane was incubated with secondary antibody (1:10000) anti-Goat IgG secondary antibody (A0545, Sigma Aldrich) in the buffer (10ml 1x TBST, 0.5g non-fat dry milk, 1 ul antibody) for 1 hour with gentle shaking at room temperature. Next, the membrane was washed 3 times, 5 min each, with 1x TBST buffer with gentle shaking. After the wash, the membrane was placed in Pierce chemiluminescence substrates for 5 minutes and imaged on film using ChemiDoc (Bio-Rad).

### PleC-CcaS chimera reporter assay

PleC-CcaS chimera reporter assays were performed based on the following steps. Starting from a -80 °C DMSO freezer stock, strains were inoculated into 5 mL LB Miller Broth in culture tubes containing appropriate antibiotics and grown at 37 °C 220 rpm for overnight. Cultures were then diluted with fresh and sterile LB media to OD_600_ =1.0 using a UV/Vis spectrophotometer (VWR, USA). The cells were then inoculated into fresh LB media with a density of 25 μl per 1mL LB media with appropriate antibiotics. The tubes were shaken at 37 °C 220 rpm until OD_600_ reaches 0.4. Then PodJ was induced with 5 mM arabinose for another 4h. After that, cells were transferred into 96-well plates. Fluorescence was measured using a 5 nm bandpass with excitation/emission for mCherry (585/nm/610nm)/CFP (456nm/480nm)/ YFP (513nm/527nm) with a manually set gain of 50. Each construct was repeated with three independent biological replicates as indicated in the standard error in the bar graph.

### Homology Modelling

PleC protein sequence was submitted into HHpred to predict protein features, fetch published crystal structures as templates and generate multiple sequences alignment^59^. A template (PDB:4GCZ) was selected to model the homology structure of PleC PAS-A and PAS-B, respectively. Homology models were then downloaded and edited with PyMol to highlight secondary structures and signal transmission motif DI/VT residues at the C-terminal of each PAS domain.

### Coupled Enzyme Activity Assay

Kinase activity was measured using a coupled enzyme assay^36^. 7.5 μM purified proteins were mixed in kinase buffer supplemented with 1.0 mM ATP, 10 mM MgCl_2_, 3 mM phosphoenolpyruvate, 0.2 mM NADH, 2 units of pyruvate kinase, and 6.6 units of lactate dehydrogenase (P0294, Sigma). Reactions were performed in three replicates in a 100 μL volume and loaded into a clear polystyrene 384 well-plate. Each reaction was initiated by adding ATP, and 340 nm absorbance was recorded every 10 s for 90 min on a Tecan M1000 microplate reader (Tecan, Switzerland). The slope of a stable, linear absorbance decay was measured to calculate ATP hydrolysis rates^60^. Background rates of ATP hydrolysis and NADH oxidation were measured and subtracted from observed ATP hydrolysis rates without adding any protein. The mean observed rate and S.D. were determined and analyzed using Prism (GraphPad).

## Acknowledgment

This research was funded through the University of Pittsburgh Start-up funds to W.S.C. We thank the Lucy Shapiro lab at Stanford University for providing the NA1000 strain used in this study. We thank Dylan Tomares and Jiefei Wang from Childers laboratory for their critiques on the experiments and writing. We thank Saumya Saurabh and Jared Schrader for discussions on biomolecular condensates in bacteria. We thank the Alex Deiters lab at the University of Pittsburgh for sharing the Tecan 100 plate reader that supported measurements in this study.

## Author contributions

W.S.C., W. Z., and C.Z. conceptualized and designed the study. C.Z. developed all the PodJ biomolecular condensate assays, PleC-CcaS chimera, and *in vitro* biochemical assays. C.Z analyzed all of the manuscript’s data. W.Z. performed and collected preliminary data on reconstituting interaction variants and implemented the final fluorescence polarization experiments and *in vitro* gel filtration experiments. K.A.K performed the initial biomolecular condensate assays. C.Z., W.Z., S.W.D., and K.A.K., constructed plasmids and strains. W.S.C. and C.Z. drafted the manuscript. All authors provided critical feedback and helped shape the research, analysis, and manuscript.

## Notes

### Competing Interest Statement

The authors have declared no competing interest.

