## Supplementary Information for "Regulation of a bacterial histidine kinase by a phase separating scaffolding protein"

**This PDF file includes:**

Figures S1 to S10
Tables S1 to S4
Text S1
SI References

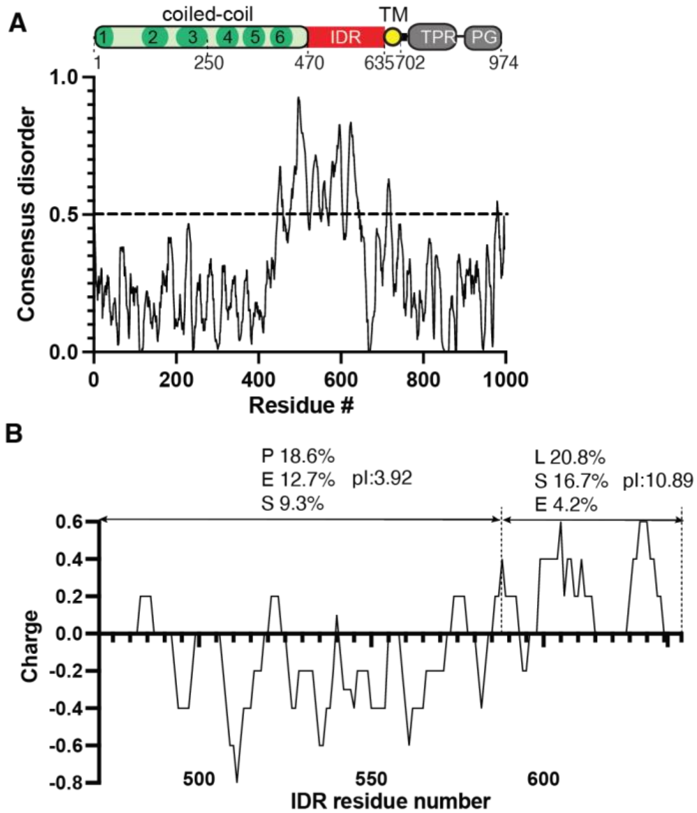

**Figure S1**. **PodJ domain architecture, IDR charge and composition analysis.** (A)PodJ domain organization predicted by HHpred and adapted from previous studies[1,2] The coiled-coil rich region was analyzed by PCOILS[3] and modeled with MODELLER[4]. The probability of intrinsic disorder over the primary sequence of PodJ represents disorder prediction from algorithm Metapredict[5]. Dash line at 0.5 on the y axis indicates the threshold for disorder probability. (B) Charge analysis of PodJ IDR in a window of 5 residues. The top 3 charged residues, their percentages and calculated pI for each block are listed.

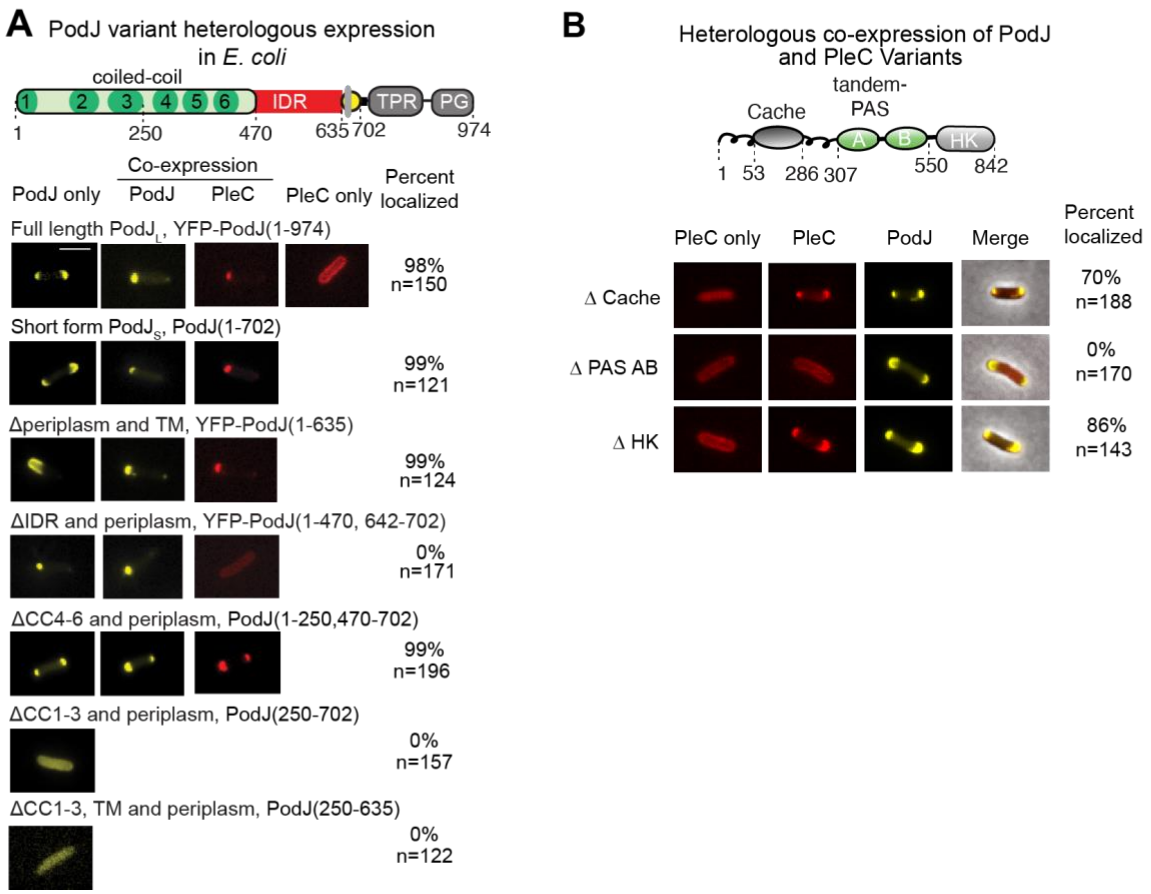

**Figure S2: Heterologous expression of PodJ and PleC variants to identify sites of interaction**. (A) Heterologous expression of a set of YFP-PodJ domain deletion variants alone or co-expressed with PleC-mcherry in *E. coli*. YFP-PodJ variants were induced with 0.5 mM IPTG, and PleC-mCherry was induced with 1mM arabinose for 2 hours. Scale bar: 2 μm. (B) Heterologous co-expression of PleC-mCherry variants with YFP-PodJ in *E. coli*. Scale bar: 2 μm. YFP-PodJ was induced with 0.5 mM IPTG, and PleC-mCherry was induced with 1 mM arabinose for 3 hours.

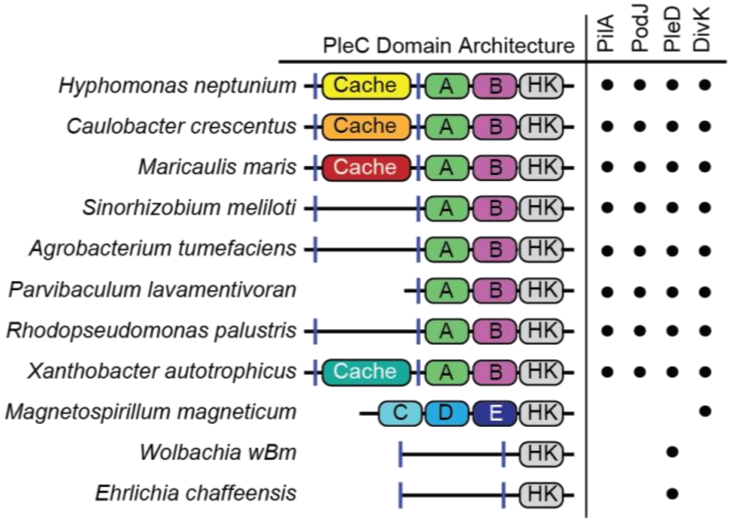

**Figure S3**. **PodJ's interaction with PleC through PAS A and B domains is conserved in other alpha-proteobacteria.** PleC domain architecture from selected alpha-proteobacteria and conservation of PleC specific inputs and cognate response regulators. Domains A and B represent both cytoplasmic PAS domains of PleC.

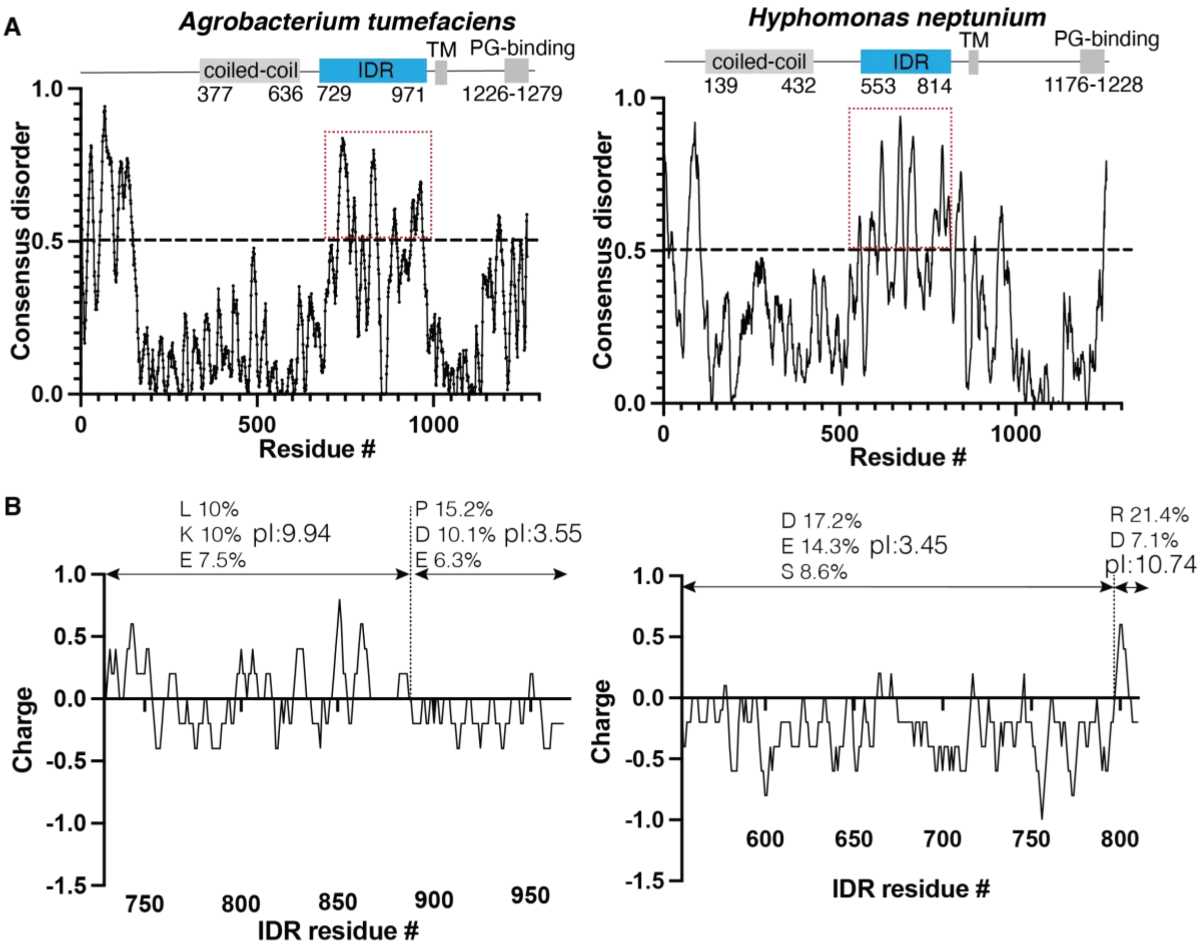

**Figure S4**. **PodJ homologs domain architecture, IDR, charge, and composition analysis from two selected alphaproteobacteria.** (A) Domain architecture and intrinsically disordered region prediction of PodJ homologs. The coiled-coil region was predicted from UniProt[6]. Transmembrane domains were predicted using TMPred[7]. Intrinsically disordered regions were predicted using metapredict[5]. Residues with disorder probability greater than 0.5 were shown in the rectangular dash red box. (B) Charge and compositional analysis of IDR in PodJ homologs. Amino acid composition and pI were calculated using EMBOSS[8]. The dominant charged residues and calculated pI for each block were also shown.

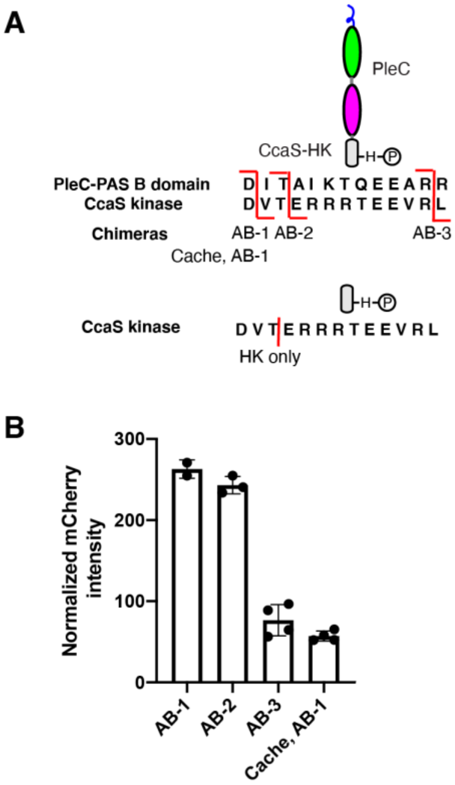

**Figure S5**. **Design and screening of PleC-CcaS chimeras that respond to PodJ expression.** (**A**). Design of PleC-CcaS chimera from light-sensing kinase CcaS and PleC. Sequence alignment between PleC PAS B and CcaS HK suggests a homology region exists after the DI/VT hinge motif. An HK-only construct without sensory domains was used as a negative control. AB indicates PleC PAS AB was used for fusion protein, while Cache AB suggests the full sensory region of PleC was used for chimera construction. (**B**). Screening of four PleC-CcaS chimeras for responsiveness to PodJ expression.

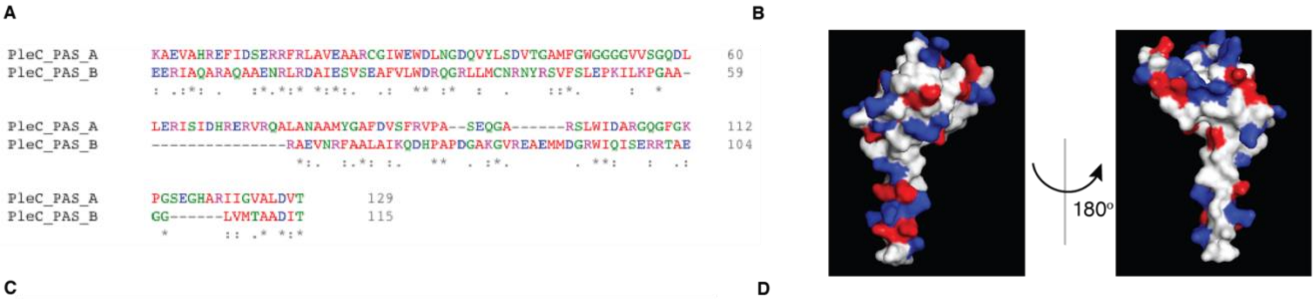

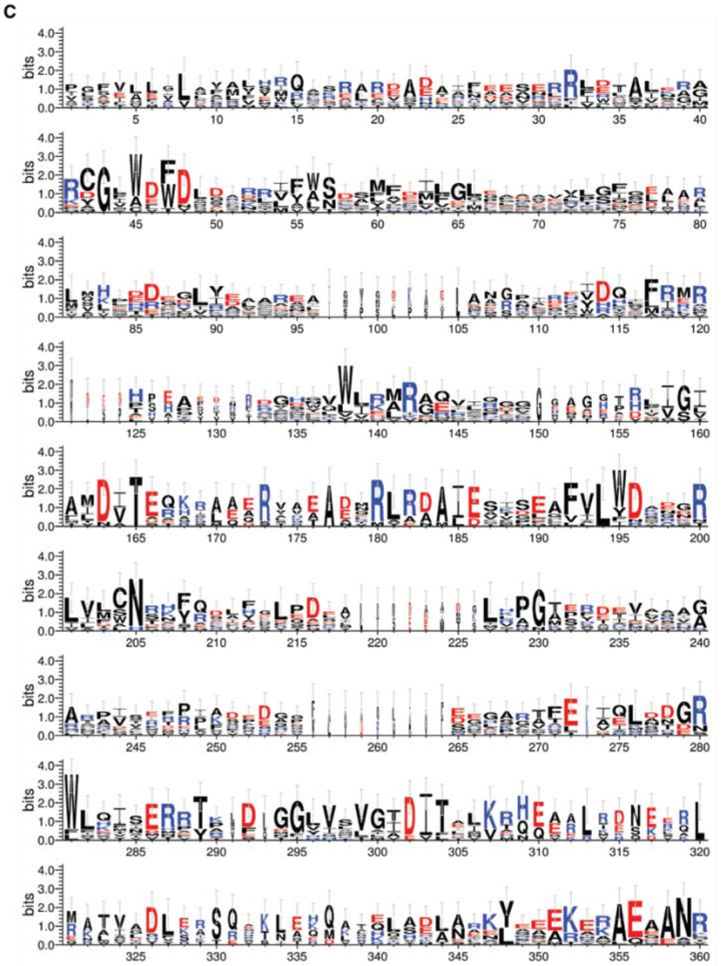

D

PAS A BLAST against alphaproteobacteria

| Accession | Per. Ident | E value |
|---|---|---|
| HEV42420.1 | 76.74 | 5.13E-56 |
| MBA4013983.1 | 73.64 | 1.29E-52 |
| MBP6878882.1 | 74.42 | 4.00E-51 |
| WP_156378276.1 | 74.42 | 4.11E-51 |
| KQW72874.1 | 74.42 | 4.34E-51 |
| WP_183774560.1 | 74.42 | 5.77E-51 |
| WP_215341570.1 | 69.77 | 8.27E-50 |
| ODU33946.1 | 73.64 | 8.96E-50 |
| THD65268.1 | 68.22 | 2.59E-49 |
| WP_056733077.1 | 73.64 | 2.74E-49 |
| MBT9472795.1 | 73.64 | 2.74E-49 |
| MBP7649181.1 | 68.99 | 5.37E-49 |
| MBP6545846.1 | 68.99 | 7.39E-49 |
| THD59163.1 | 66.67 | 8.72E-49 |
| RYF94744.1 | 75.21 | 2.00E-48 |
| THD79946.1 | 65.89 | 1.38E-47 |
| WP_012523228.1 | 65.12 | 2.03E-47 |
| MBU2134991.1 | 72.87 | 5.25E-47 |
| WP_111513056.1 | 65.89 | 8.72E-47 |
| MBI5942540.1 | 71.32 | 2.17E-46 |
| MAK81792.1 | 72.87 | 4.00E-46 |
| WP_111456050.1 | 64.34 | 1.07E-45 |
| WP_111530019.1 | 66.14 | 2.69E-45 |
| OYX31829.1 | 61.60 | 3.57E-43 |
| OYU70959.1 | 66.67 | 4.38E-43 |
| MBI1196418.1 | 64.71 | 6.66E-43 |
| MBA4792159.1 | 68.99 | 1.07E-42 |
| MBU1376023.1 | 62.50 | 3.54E-42 |
| MBL8773636.1 | 63.03 | 4.24E-42 |
| PZQ63279.1 | 61.67 | 4.99E-42 |
| WP_111276259.1 | 63.03 | 6.35E-42 |
| OXE37280.1 | 67.44 | 1.26E-41 |
| WP_156466929.1 | 61.67 | 2.33E-41 |
| TAJ70067.1 | 62.19 | 2.92E-41 |
| MBJ7411368.1 | 61.67 | 2.96E-41 |
| OHB27668.1 | 61.67 | 3.28E-41 |
| Average | **68.28** | |

PAS B BLAST against alphaproteobacteria

| Accession | Per. Ident | E value | | Accession | Per. Ident | E value |
|---|---|---|---|---|---|---|
| HEV42420.1 | 91.30 | 3.47E-58 | | PZU56446.1 | 57.39 | 1.65E-35 |
| OYW90456.1 | 85.22 | 3.88E-54 | | WP_207870595.1 | 54.78 | 1.66E-35 |
| MBT9472795.1 | 85.22 | 1.01E-53 | | EGF96315.1 | 54.78 | 2.61E-35 |
| WP_056733077.1 | 85.22 | 1.19E-53 | | EDK81457.1 | 56.52 | 2.63E-35 |
| RYF94744.1 | 85.22 | 1.24E-53 | | PZO08317.1 | 56.52 | 2.82E-35 |
| MAK81792.1 | 86.09 | 1.23E-52 | | WP_081836168.1 | 56.52 | 2.82E-35 |
| MBI5942540.1 | 81.74 | 1.88E-51 | | WP_183196426.1 | 56.52 | 2.89E-35 |
| MBA4013983.1 | 82.61 | 1.99E-51 | | MBB3872164.1 | 56.52 | 2.93E-35 |
| WP_183774560.1 | 82.61 | 1.99E-51 | | OGN47325.1 | 56.52 | 3.26E-35 |
| WP_156378276.1 | 82.61 | 2.25E-51 | | MBJ7317341.1 | 56.52 | 3.34E-35 |
| KQW72874.1 | 82.61 | 2.30E-51 | | KDP94631.1 | 56.52 | 3.36E-35 |
| MBP7649181.1 | 83.48 | 2.41E-51 | | WP_040349769.1 | 56.52 | 3.36E-35 |
| MBP6545846.1 | 83.48 | 3.52E-51 | | OGN48782.1 | 56.52 | 3.36E-35 |
| ODU33946.1 | 82.61 | 4.09E-51 | | MBU1273609.1 | 56.52 | 3.41E-35 |
| TAJ70067.1 | 73.04 | 5.18E-51 | | WP_196066342.1 | 56.52 | 3.46E-35 |
| WP_111530019.1 | 73.04 | 6.81E-51 | | RYG16086.1 | 53.04 | 3.51E-35 |
| WP_215341570.1 | 82.61 | 9.66E-51 | | HAF79465.1 | 54.37 | 3.69E-35 |
| MBL8554656.1 | 70.44 | 5.61E-50 | | TAJ56292.1 | 54.78 | 4.50E-35 |
| PZQ63279.1 | 72.17 | 7.69E-50 | | WP_090647314.1 | 56.52 | 5.21E-35 |
| MBP6878882.1 | 80.87 | 8.37E-50 | | SCW53406.1 | 56.52 | 6.21E-35 |
| WP_111276259.1 | 71.30 | 1.06E-49 | | MBI2261230.1 | 55.65 | 7.03E-35 |
| MBJ7411368.1 | 72.17 | 2.13E-49 | | MBA3049353.1 | 54.78 | 1.12E-34 |
| MBU1376023.1 | 71.30 | 2.18E-49 | | WP_107871417.1 | 56.52 | 1.23E-34 |
| OHB27668.1 | 72.17 | 3.29E-49 | | WP_013479453.1 | 58.26 | 1.26E-34 |
| WP_156466929.1 | 71.30 | 3.44E-49 | | WP_126421283.1 | 58.26 | 1.27E-34 |
| MBL8773636.1 | 70.44 | 1.05E-48 | | BBF80745.1 | 58.26 | 1.27E-34 |
| MBA4792159.1 | 81.74 | 1.50E-48 | | WP_183203771.1 | 55.65 | 1.48E-34 |
| WP_012523228.1 | 71.30 | 1.15E-47 | | RZJ29588.1 | 53.15 | 1.92E-34 |
| MBQ1542868.1 | 79.13 | 1.52E-47 | | WP_123287820.1 | 56.52 | 2.60E-34 |
| WP_111456050.1 | 73.04 | 3.01E-47 | | WP_040344694.1 | 54.78 | 3.32E-34 |
| OYX31829.1 | 71.30 | 4.09E-47 | | OGN50218.1 | 55.65 | 3.82E-34 |
| THD59163.1 | 78.26 | 4.66E-47 | | MBB4082137.1 | 55.65 | 4.09E-34 |
| ODT85277.1 | 69.57 | 8.90E-47 | | WP_087140987.1 | 54.78 | 4.30E-34 |
| THD65268.1 | 75.65 | 2.14E-45 | | HAV50066.1 | 56.52 | 4.31E-34 |
| MBI1196418.1 | 67.83 | 1.07E-44 | | MAL88326.1 | 56.52 | 4.62E-34 |
| OYU70959.1 | 73.04 | 8.39E-42 | | OYK21121.1 | 54.78 | 5.23E-34 |
| THD79946.1 | 63.48 | 1.33E-40 | | WP_076223497.1 | 54.78 | 5.88E-34 |
| MBE7219139.1 | 64.91 | 3.07E-39 | | WP_128719619.1 | 54.78 | 5.88E-34 |
| WP_168045073.1 | 59.13 | 1.46E-38 | | MBD3817583.1 | 54.78 | 5.99E-34 |
| WP_111513056.1 | 68.70 | 2.91E-38 | | WP_088491656.1 | 54.78 | 6.17E-34 |
| MBF0665016.1 | 60.00 | 7.01E-38 | | WP_088411220.1 | 54.78 | 6.23E-34 |
| WP_029087896.1 | 60.87 | 2.01E-37 | | WP_128115884.1 | 54.78 | 7.28E-34 |
| MBJ7485572.1 | 57.39 | 2.28E-37 | | ESQ86095.1 | 56.52 | 9.96E-34 |
| NEW09018.1 | 55.65 | 2.21E-36 | | WP_081714526.1 | 56.52 | 1.12E-33 |
| MBU2271017.1 | 56.52 | 2.53E-36 | | WP_178827671.1 | 54.78 | 1.33E-33 |
| MBU1383432.1 | 56.52 | 2.73E-36 | | WP_114817891.1 | 54.78 | 1.39E-33 |
| MBU2417434.1 | 56.52 | 2.99E-36 | | MBD3833676.1 | 54.78 | 1.50E-33 |
| WP_183254730.1 | 57.39 | 5.87E-36 | | WP_141628378.1 | 53.04 | 1.60E-33 |
| MBN8552289.1 | 59.13 | 7.85E-36 | | WP_023460854.1 | 55.65 | 1.71E-33 |
| OXE37280.1 | 68.70 | 1.14E-35 | | | | |
| MBD3835210.1 | 57.39 | 1.16E-35 | | Average | **64.44** | |

**Figure S6**. **Bioinformatic analysis of PleC.** (**A**). Sequence similarity analysis of alignment PleC PAS A and B domains (**B**). Illustration of surface-exposed charged residues of PleC PAS B domain homology model. Color code: white (neutral), red (negative) and blue (positive). (**C**). Sequence logo[9] of PleC PAS AB domain from residue 290-580. (D) Sequence similarity analysis of PleC PAS A and B domains. PleC PAS A and B domains were searched in BLAST against alphaproteobacteria, and the average sequence similarity was calculated for each domain.

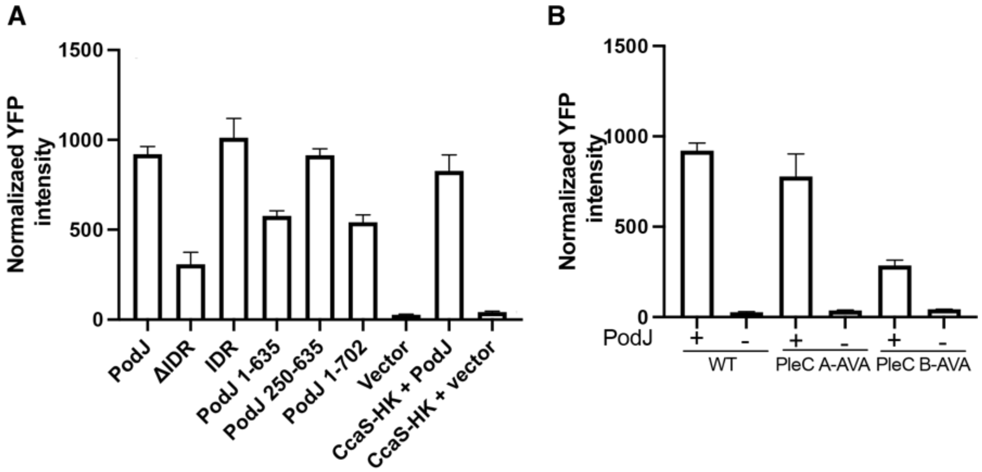

**Figure S7. Quantification of YFP-PodJ and its variant expression level in PleC-CcaS reporter system assays in *E. coli.*** (A). The expression level of YFP-PodJ variants (B). The expression level of YFP-PodJ in PleC PAS A and B signal transmission DI/VT motif and mutants.

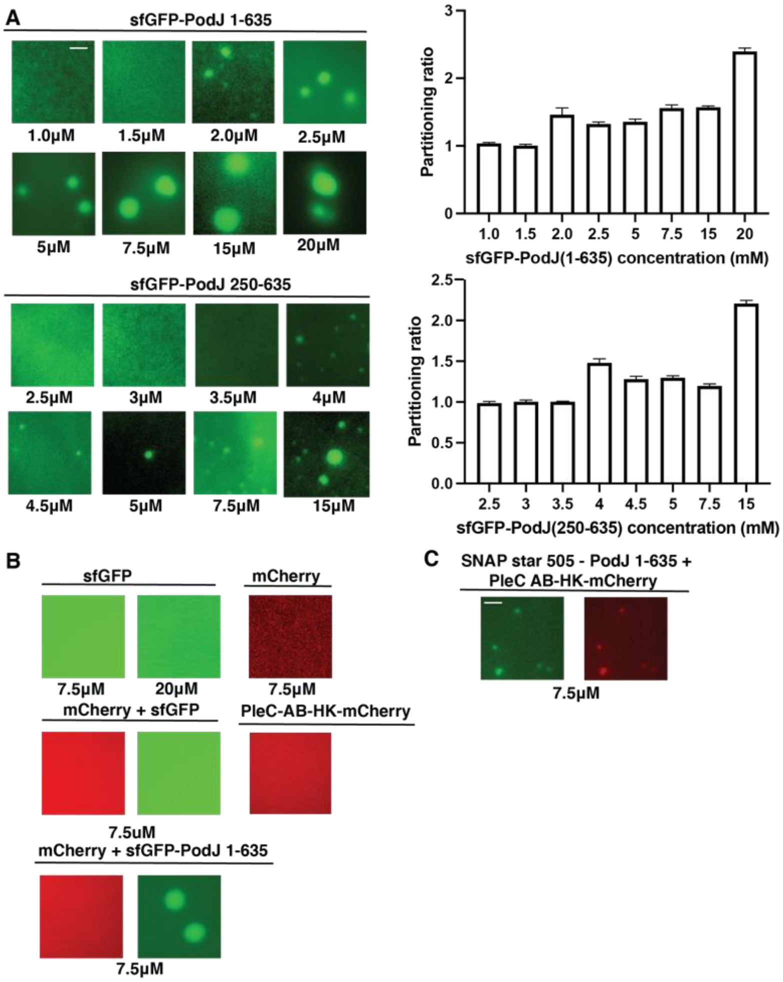

**Figure S8**. **PodJ's ability to form biomolecular condensates depends on coiled-coil 1-3 region but not fluorescent protein.** (A). Concentration dependency of sfGFP-PodJ1-635 and sfGFP-PodJ250-635 phase separation. (B) PodJ recruits PleC into biomolecular condensates independent of fluorescent proteins. sfGFP, mCherry, mCherry plus sfGFP did not phase separate at low PEG8000 concentrations. PleC AB-HK only forms biomolecular condensates at relatively high PEG8000 concentrations. mCherry was not recruited by sfGFP-PodJ 1-635 at various PEG 8000 concentrations. (C) Swapping sfGFP to SNAP-tag retained PodJ's droplet formation as well as PleC AB-HK-mCherry recruitment capability. All samples were prepared in 50 mM Tris-HCl, pH=8.0, 200mM KCl with varying amounts of proteins unless otherwise specified at room temperature. Representative images were shown from the whole field of imaging. Scale bar: 2.5 µm.

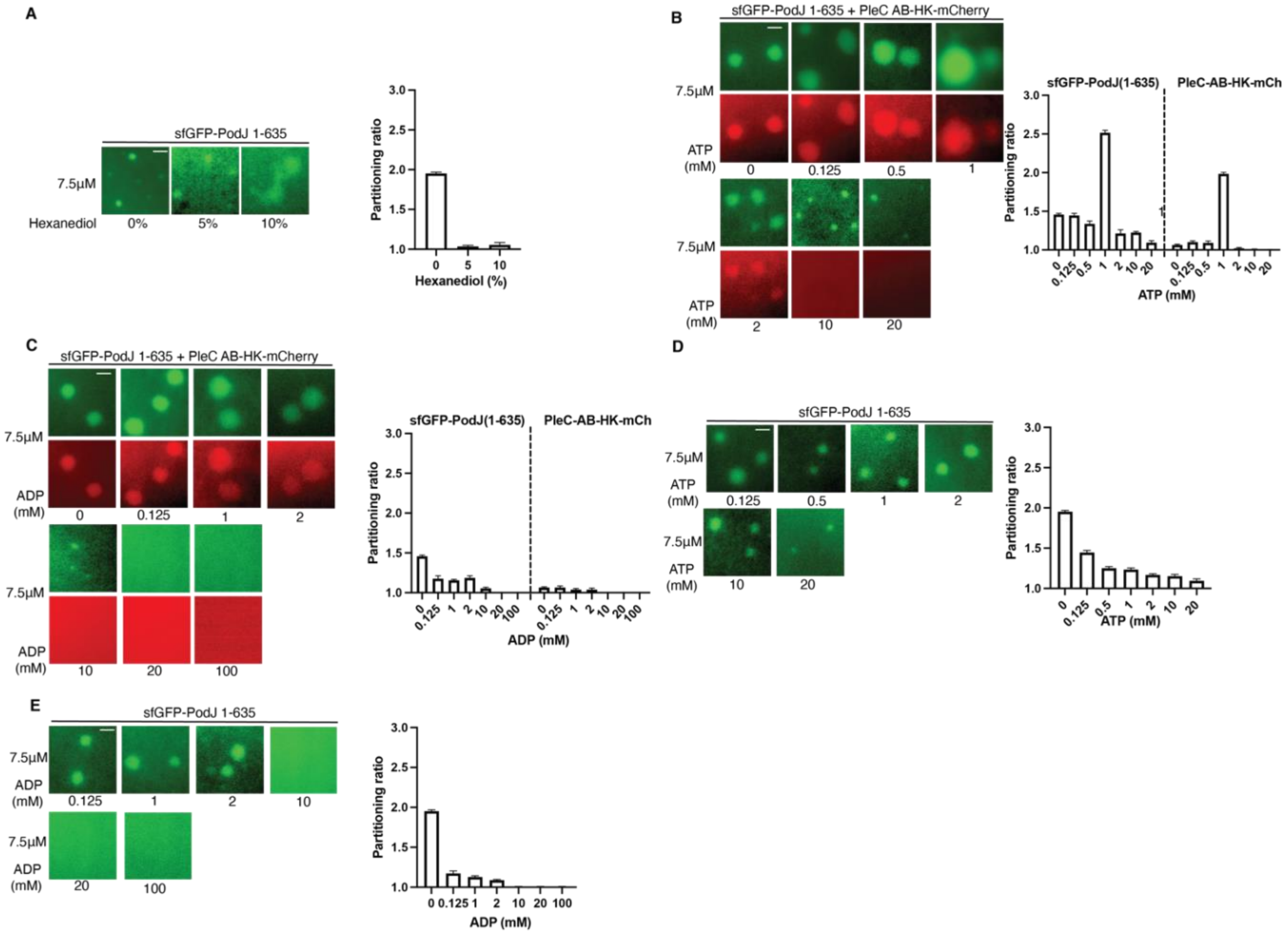

**Figure S9. PodJ biomolecular condensates material properties are regulated by hexanediol, ATP or ADP addition *in vitro*.** (A). Hexanediol disrupts well-organized droplet formation of sfGFP-PodJ 1-635 in a dose-dependent manner. (B). The addition of ATP regulates droplet formation and dissolution of the sfGFP-PodJ 1-635 and PleC AB-HK-mCherry complex in the droplets in a dose-dependent manner. (C) The addition of ADP regulates droplet formation and dissolution of the sfGFP-PodJ 1-635 and PleC AB-HK-mCherry complex in the droplets in a dose-dependent manner. (D) ATP acts as a hydrotrope that leads to the dissolution of droplets at high concentrations. (E) ADP acts as a hydrotrope that leads to the dissolution of droplets at high concentrations.

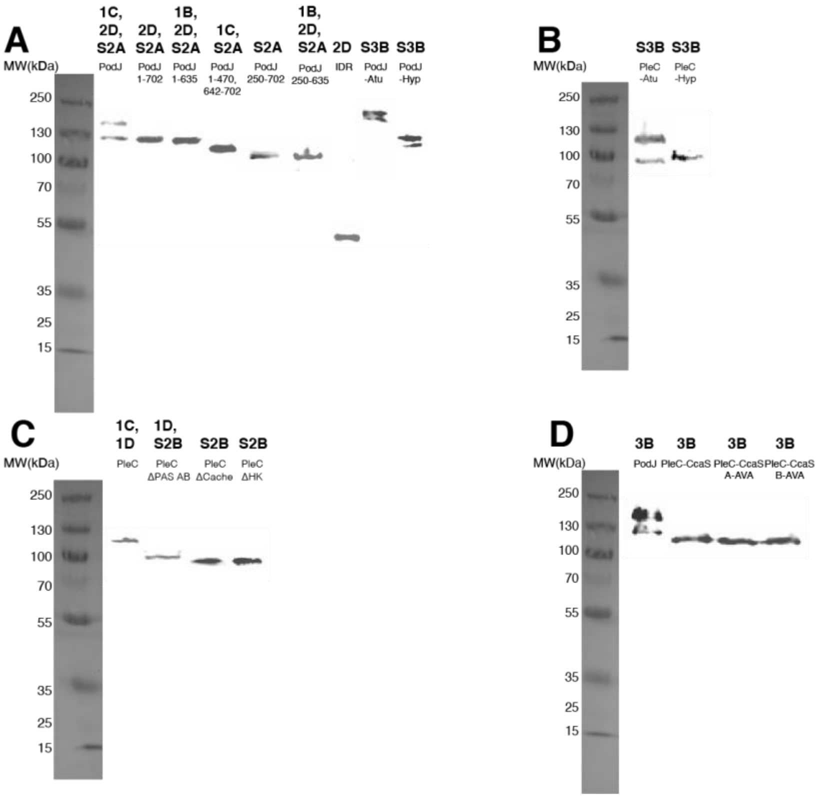

**Figure S10 Western blotting for constructs used in this work**. (A). YFP/sfGFP-PodJ variants from *Caulobacter* expressed in *E coli* or *Caulobacter* and from alphaproteobacteria species expressed in *E coli*. (B). CFP-PleC from alphaproteobacteria species expressed in *E coli*. (C). PleC-mCherry variants from *Caulobacter* expressed in *E coli* or *Caulobacter.* (D). YFP-PleC-CcaS signal transmission motif expressed in *E coli.* The figure that each construct corresponds to is listed on top of each construct name in bold. Western blot analysis indicates each construct was expressed and exhibited little proteolysis. The exception is that full-length PodJ constructs, even when expressed in *E. coli*, exhibit proteolysis into a shorter form of PodJ.

### Table S1. List of PCR reactions for Gibson assembly

| Plasmid | First Insert Forward Primer | First Insert Reverse Primer | Second Insert Forward Primer | Second Insert Reverse Primer | Third Insert Forward Primer | Third Insert Reverse Primer |
|---|---|---|---|---|---|---|
| pWZ21 | WZ0044_(Plec)_forward | WZ0045_(Plec)_reverse | WZ0046_(mcherry)_forward | WZ0024_(mcherry)_reverse | | |
| pWZ24 | WZ0050_(PBAD)_forward | WZ0051_(PBAD)_reverse | WZ0052_(YFP-PODJ)_forward | WZ0053_(YFP-PODJ)_reverse | | |
| pWZ12 | WZ0025_(pxyfpc-6)_forward | WZ0010_(PCDF-YFP)_reverse | WZ0011_(HRSAT)_(PODJ)_forward | WZ0026_(PodJ)_reverse | | |
| pWZ138-kan | WZ0324_(pbvmcs-6)_forward | WZ0325_(pbvmcs-6)_reverse | WZ0326_(plec-mch)_forward | WZ0327_(plec-mch)_reverse | | |
| pWZ138-chlor | WZ0324_(pbvmcs-6)_forward | WZ0325_(pbvmcs-6)_reverse | WZ0326_(plec-mch)_forward | WZ0327_(plec-mch)_reverse | | |
| pCZ340 | CZ10807_(pCDF)_forward | CZ10808_(pCDF)_reverse | CZ10809_(Atu_PleC)_forward | CZ10810_(Atu_PleC)_reverse | | |
| pCZ341 | CZ10811_(pCDF)_forward | CZ10812_(pCDF)_reverse | CZ10813_(Xan_PleC)_forward | CZ10814_(Xan_PleC)_reverse | | |
| pCZ342 | CZ10815_(pCDF)_forward | CZ10816_(pCDF)_reverse | CZ10817_(Hyp_PleC)_forward | CZ10818_(Hyp_PleC)_reverse | | |
| pWZ127 | WZ0173_(pbad)_forward | WZ0312_(pbad)_reverse | WZ0315_(plec-pasc)_forward | WZ0316_(plec-pasc)_reverse | WZ0317_(mcherry)_forward | WZ0176_(plec-mcherry)_reverse |
| pWZ135 | WZ0173_(pbad)_forward | WZ0322_(pbad)_reverse | WZ0323_(plec-pasd-atg)_forward | WZ0320_(plec-pasd)_reverse | WZ0321_(mcherry)_forward | WZ0176_(plec-mcherry)_reverse |
| pWZ136 | WZ0173_(pbad)_forward | WZ0312_(pbad)_reverse | WZ0315_(plec-pasc)_forward | WZ0320_(plec-pasd)_reverse | WZ0321_(mcherry)_forward | WZ0176_(plec-mcherry)_reverse |
| YFP-J23102-pCZ267 | CZ10625_(pCZ10224)_forward | CZ10622_(Delta_CcaS)_reverse | CZ10623_(eyfp)_forward | CZ10626_(eyfp)_reverse | | |
| pCZ259 | CZ10596_(pCZ130)_forward | CZ10597_(pCZ130)_reverse | CZ10598_(CloDF13)_forward | CZ10599_(CloDF13)_reverse | | |
| pCZ253 | CZ10583_(CcaS_with_TM)_forward | CZ10524_(pSR43_6)_reverse | CZ10525_(PleC_PAS3)_forward | CZ10584_(PleC_CD)_reverse | | |
| pW173 | WZ0050_(PBAD)_forward | WZ0186_(pcdf-mcherry)_reverse | WZ0187_(PODJ1-470)_forward | WZ0446_(podjdeltaPSE)_reverse | | |
| pCZ412 | CZ11031_(pBAD-YFP)_forward | CZ11032_(pBAD-YFP)_reverse | CZ11033_(PSE)_forward | CZ11034_(PSE)_reverse | | |
| pCZ253 (D433A, T435A) | CZ-pCZ253-DVT-C-AVA-F | CZ-pCZ253-DVT-C-AVA-R | | | | |
| pCZ253 (D548A, I549V, T550A) | CZ11057_(pWZ21)_forward | CZ11058_(pWZ21)_reverse | CZ11059_(PleC_CD_D_DVT)_forward | CZ11060_(PleC_CD_D_DVT)_reverse | | |
| pCZ253 (D433A, T435A, D548A, I549V, T550A) | CZ11057_(pWZ21)_forward | CZ11058_(pWZ21)_reverse | CZ11059_(PleC_CD_D_DVT)_forward | CZ11060_(PleC_CD_D_DVT)_reverse | | |
| pCZ391 | CZ10973_(pTEV5)_forward | CZ10974_(pTEV5)_reverse | CZ10975_(msfGFP)_forward | CZ10976_(msfGFP)_(stop)_reverse | | |
| pmas00001 | mas00001_(pTEV5)_forward | mas00002_(pTEV5)_reverse | mas00003_(mCherry)_forward | mas00004_(mCherry)_reverse | | |
| pCZ388 | CZ10944_(pTEV5-bb)_forward | CZ10964_(pTEV5-bb)_(S)_reverse | CZ10965_(msfGFP)_forward | CZ10966_(msfGFP)_(HRSAT)_reverse | CZ10948_(PodJ-1-635)_forward | CZ10949_(PodJ-1-635)_reverse |
| pmas00002 | mas00001_(pTEV5)_forward | mas00002_(pTEV5)_reverse | mas00003_(pWZ21)_forward | mas00004_(pWZ21)_reverse | | |
| pCZ388-dCC-13 | CZ-msfGFP-dCC-1-3-F | CZ-msfGFP-dCC-1-3-R | | | | |
| pCZ382 | CZ10944_(pTEV5-bb)_forward | CZ10945_(pTEV5-bb)_reverse | CZ10946_(SNAP)_forward | CZ10947_(SNAP)_(HRSAT)_reverse | CZ10948_(PodJ-1-635)_forward | CZ10949_(PodJ-1-635)_reverse |
| pCZ479 | pWZ24-dCC1-3-F1 | pWZ24-dCC1-3-R1 | | | | |
| WZ365 | WZH566-pxyfpn-2-Pxyl-sfGFP-PodJ1-635-F | WZH567-pxyfpn-2-Pxyl-sfGFP-PodJ1-635-R | | | | |
| pSWD259 | SWD700_(start)_(sfgfp)_forward | SWD790_(sfgfp)_(HRSAT)_reverse | SWD791_(podJ_251_670)_forward | SWD792_(podJ_251_670)_(stop)_reverse | | |

### Table S2. List of PCR primers

| Name | Description |
|---|---|
| CZ10524_(pSR43_6)_reverse | GCGGCTCTGGATCATTAAAGATTGGCGGATATGTTGGG |
| CZ10525_(PleC_PAS3)_forward | CCGCCAATCTTTAATGATCCAGAGCCGCAAGGCCG |
| CZ10583_(CcaS_with_TM)_forward | GACCGCCGCCGACGTCACTGAGCGCCGACGC |
| CZ10584_(PleC_CD)_reverse | GGCGCTCAGTGACGTCGGCGGCGGTCATGACAAGACC |
| CZ10596_(pCZ130)_forward | GCAATTTATCTCTTCAAATGTACTAGTGCTTGGATTCTCACCAA |
| CZ10597_(pCZ130)_reverse | GCTATTTGTTTATTTTTCTTTTTACGGTTCCTGGCCTTTTGC |
| CZ10598_(CloDF13)_forward | GGCCAGGAACCGTAAAAAGAAAAATAAACAAATAGCTAGCTCACTCGGTCG |
| CZ10599_(CloDF13)_reverse | GGTGAGAATCCAAGCACTAGTACATTTGAAGAGATAAATTGCACTGAAATC |
| CZ10622_(Delta_CcaS)_reverse | CCTTGCTCACCATTGCGCCTTCCTCCTATTAAGATTTTAACC |
| CZ10623_(eyfp)_forward | GGAGGAAGGCGCAATGGTGAGCAAGGGCGAGGAGC |
| CZ10625_(pCZ10224)_forward | CCGGTCGGCCACCATGATCCAGAGCCGCAAGGC |
| CZ10626_(eyfp)_reverse | GGCTCTGGATCATGGTGGCCGACCGGTGCTTGTACAGC |
| CZ10807_(pCDF)_forward | GGTGTGTGGGACGGTGCATCACCGGTCGGCCACCATGG |
| CZ10808_(pCDF)_reverse | GCACGTTCGTCATCTGGCTGTGGTGATGATGGTGATGGC |
| CZ10809_(Atu_PleC)_forward | TCACCACAGCCAGATGACGAACGTGCGGCGGG |
| CZ10810_(Atu_PleC)_reverse | TGGCCGACCGGTGATGCACCGTCCCACACACCCC |
| CZ10811_(pCDF)_forward | GGCCGCGGTGATGCACCGGTCGGCCACCATGG |
| CZ10812_(pCDF)_reverse | CGTTGGCGCGTGCCATCTGGCTGTGGTGATGATGGTGATGGC |
| CZ10813_(Xan_PleC)_forward | TCACCACAGCCAGATGGCACGCGCCAACGCT |
| CZ10814_(Xan_PleC)_reverse | TGGCCGACCGGTGCATCACCGCGGCCAGCCT |
| CZ10815_(pCDF)_forward | GCGGGATGTTGCCCACCGGTCGGCCACCATGG |
| CZ10816_(pCDF)_reverse | CCCCGCGAATCATCTGGCTGTGGTGATGATGGTGATGGC |
| CZ10817_(Hyp_PleC)_forward | TCACCACAGCCAGATGATTCGCGGGGAAAAGGTAAGGC |
| CZ10818_(Hyp_PleC)_reverse | TGGCCGACCGGTGGGCAACATCCCGCTGGATGTCG |
| CZ11031_(pBAD-YFP)_forward | AAGGCGCGCTTGTAAGAAGCTTGGCTGTTTTGGCGG |
| CZ11032_(pBAD-YFP)_reverse | GACGGGGTGGCCGACCGGTGCTTG |
| CZ11033_(PSE)_forward | TACAAGCACCGGTCGGCCACCCCGTCGCCGCCGCCCGCG |
| CZ11034_(PSE)_reverse | AAAACAGCCAAGCTTCTTACAAGCGCGCCTTCGACTTCTTGG |
| CZ11057_(pWZ21)_forward | GACCGCCGCCGACATCACGGCCATCAAGACCCAGG |
| CZ11058_(pWZ21)_reverse | GGCTCTGGATCATCAGCAGCAGCGCCAGGGC |
| CZ11059_(PleC_CD_D_DVT)_forward | GGCGCTGCTGCTGATGATCCAGAGCCGCAAGGCCG |
| CZ11060_(PleC_CD_D_DVT)_reverse | GGTCTTGATGGCCGTGATGTCGGCGGCGGTCATGACAAGACC |
| RecUni-1 | ATGCCGTTTGTGATGGCTTCCATGTCG |
| RecXyl-2 | TCTTCCGGCAGGAATTCACTCACGCC |
| WZ0010_(PCDF-YFP)_reverse | GGTGGCCGACCGGTGCTTGTACAGCTCGTCCATGCCG |
| WZ0011_(HRSAT)_(PODJ)_forward | GACGAGCTGTACAAGCACCGGTCGGCCACCATGACGGCGGCTTCGCCATGG |
| WZ0024_(mcherry)_reverse | GTCGACCTGCAGGCGCGCCGAGCTCGAATTCTTACTTGTACAGCTCGTCCATGCCGC |

| | |
|---|---|
| WZ0025_(pxyfpc-6)_forward | CTTAGTATATTAGTTAAGTATAAGAAGGAGATATAATGGTGAGCAAGGGCGAGGAGC |
| WZ0026_(PodJ)_reverse | GTGGCCGGCCGATATCCAATTGAGATCTGCTTAGCGCGCGTAGACCGACAGG |
| WZ0044_(Plec)_forward | CAGCAGCCATCACCATCATCACCACAGCCAATGGGCAGACACGGGGGGC |
| WZ0045_(Plec)_reverse | GGTGGCCGACCGGTGGGCCGCCACGAAGTCGCG |
| WZ0046_(mcherry)_forward | GACTTCGTGGCGGCCCACCGGTCGGCCACCATGG |
| WZ0050_(PBAD)_forward | GGTCTACGCGCGCTAAGAAGCTTGGCTGTTTTGGCGG |
| WZ0050_(PBAD)_forward | GGTCTACGCGCGCTAAGAAGCTTGGCTGTTTTGGCGG |
| WZ0051_(PBAD)_reverse | CGCCCTTGCTCACCATTTAATTCCTCCTGTTAGCCCAAAAA |
| WZ0052_(YFP-PODJ)_forward | AACAGGAGGAATTAAATGGTGAGCAAGGGCGAGGAGC |
| WZ0053_(YFP-PODJ)_reverse | AAACAGCCAAGCTTCTTAGCGCGCGTAGACCGACAGG |
| WZ0173_(pbad)_forward | CGAGCTGTACAAGTAAGAAGCTTGGCTGTTTTGGCGG |
| WZ0176_(plec-mcherry)_reverse | CCAAAACAGCCAAGCTTCTTACTTGTACAGCTCGTCCATGCCG |
| WZ0186_(pcdf-mcherry)_reverse | CGAAGCCGCCGTCATGGTGGCCGACCGGTGCTTGTA |
| WZ0187_(PODJ1-470)_forward | CACCGGTCGGCCACCATGACGGCGGCTTCGCCATGG |
| WZ0312_(pbad)_reverse | GCGGCTCTGGATCATTTAATTCCTCCTGTTAGCCCAAAAA |
| WZ0315_(plec-pasc)_forward | CTAACAGGAGGAATTAAATGATCCAGAGCCGCAAGGCCG |
| WZ0316_(plec-pasc)_reverse | GGTGGCCGACCGGTGGGTGACGTCCAGCGCCACGC |
| WZ0317_(mcherry)_forward | GCGCTGGACGTCACCCACCGGTCGGCCACCATGGTGAGC |
| WZ0320_(plec-pasd)_reverse | GGTGGCCGACCGGTGCGTGATGTCGGCGGCGGTCATGACAAGACC |
| WZ0321_(mcherry)_forward | GCCGCCGACATCACGCACCGGTCGGCCACCATGGTGAGC |
| WZ0322_(pbad)_reverse | CGATCCGCTCCTCCATTTAATTCCTCCTGTTAGCCCAAAAA |
| WZ0323_(plec-pasd-atg)_forward | GCTAACAGGAGGAATTAAATGGAGGAGCGGATCGCCCAGG |
| WZ0324_(pbvmcs-6)_forward | CGAGCTGTACAAGTAAAAAACGGGCCCCCCCTCGAGG |
| WZ0325_(pbvmcs-6)_reverse | CCCGTGTCTGCCCATCGTTTCCTCGCATCGTGGTTCGG |
| WZ0326_(plec-mch)_forward | CGATGCGAGGAAACGATGGGCAGACACGGGGGGCC |
| WZ0327_(plec-mch)_reverse | GGGGGGGCCCGTTTTTTACTTGTACAGCTCGTCCATGCCG |
| WZ0446_(podjdeltaPSE)_reverse | CAAAACAGCCAAGCTTCTTAGCGCGCGTAGACCGACAGG |
| CZ-pCZ253-DVT-C-AVA-F | TCGGCGTGGCGCTGGCGGTCGCGGAGGAGCGGATCGCCCAGGCCC |
| CZ-pCZ253-DVT-C-AVA-R | GACCGCCAGCGCCACGCCGATGATGCGGGCGTGAC |
| CZ10973_(pTEV5)_forward | GAACTCTACAAATAGTAAGCTAGCCATATGGCCATGGA |
| CZ10974_(pTEV5)_reverse | CTTCACCTTTAGACATGCCCTGAAAATACAGGTTTTCACTAGTTGGG |
| CZ10975_(msfGFP)_forward | GTATTTTCAGGGCATGTCTAAAGGTGAAGAACTGTTCACCGGTGTTGT |
| CZ10976_(msfGFP)_(stop)_reverse | GGCCATATGGCTAGCTTACTATTTGTAGAGTTCATCCATGCCGTGCGTG |
| mas00001_(pTEV5)_forward | CGAGCTGTACAAGTAATAGGCTAGCCATATGGCCATGGA |
| mas00002_(pTEV5)_reverse | GCCCTTGCTCACCATGCCCTGAAAATACAGGTTTTCACTAGTTGGG |
| mas00003_(mCherry)_forward | GTATTTTCAGGGCATGGTGAGCAAGGGCGAGGAGG |
| mas00004_(mCherry)_reverse | CCATATGGCTAGCCTATTACTTGTACAGCTCGTCCATGCCGC |
| CZ10944_(pTEV5-bb)_forward | CGAAGGCGCGCTTGTAAGCTAGCCATATGGCCATGGA |
| CZ10964_(pTEV5-bb)_(S)_reverse | GGTGAACAGTTCTTCACCTTTAGACATGCCCTGAAAATACAGGTTTTCACTAGTTGGG |
| CZ10965_(msfGFP)_forward | TCAGGGCATGTCTAAAGGTGAAGAACTGTTCACCGGTGTTGT |
| CZ10966_(msfGFP)_(HRSAT)_reverse | CGAAGCCGCCGTCATGGTGGCCGACCGGTGTTTGTAGAGTTCATCCATGCCGTGCGT |

| | |
|---|---|
| CZ10948_(PodJ-1-635)_forward | CCGGTCGGCCACCATGACGGCGGCTTCGCCA |
| CZ10949_(PodJ-1-635)_reverse | TATGGCTAGCTTACAAGCGCGCCTTCGACTTCTTGGT |
| mas00001_(pTEV5)_forward | GGACGAGCTGTACAAGTAGGCTAGCCATATGGCCATGGA |
| mas00002_(pTEV5)_reverse | CGACCTCGGCCTTGCCCTGAAAATACAGGTTTTCACTAGTTGGG |
| mas00003_(pWZ21)_forward | CCTGTATTTTCAGGGCAAGGCCGAGGTCGCCCATCG |
| mas00004_(pWZ21)_reverse | GCCATATGGCTAGCCTACTTGTACAGCTCGTCCATGCCGCC |
| CZ-msfGFP-dCC-1-3-F | ACAAACACCGGTCGGCCACCTTGGGCGCCGTCGAGACTGCCAATC |
| CZ-msfGFP-dCC-1-3-R | GGTGGCCGACCGGTGTTTGTAGAGTTCATCCATGCCGTGCGT |
| CZ10944_(pTEV5-bb)_forward | CGAAGGCGCGCTTGTAAGCTAGCCATATGGCCATGGA |
| CZ10945_(pTEV5-bb)_reverse | CGCAATCTTTGTCCATGCCCTGAAAATACAGGTTTTCACTAGTTGGG |
| CZ10946_(SNAP)_forward | TTTTCAGGGCATGGACAAAGATTGCGAAATGAAACGTACCACCC |
| CZ10947_(SNAP)_(HRSAT)_reverse | CGAAGCCGCCGTCATGGTGGCCGACCGGTGGGATCCTGGCGCGCCTATACC |
| CZ10948_(PodJ-1-635)_forward | CCGGTCGGCCACCATGACGGCGGCTTCGCCA |
| CZ10949_(PodJ-1-635)_reverse | TATGGCTAGCTTACAAGCGCGCCTTCGACTTCTTGGT |
| pWZ24-dCC1-3-F1 | TTGGGCGCCGTCGAGACTGCCAATCCCGCCACGGG |
| pWZ24-dCC1-3-R1 | GGCAGTCTCGACGGCGCCCAAGGTGGCCGACCGGTGCTTGTACAG |
| WZH566-pxyfpn-2-Pxyl-sfGFP-PodJ1-635-F | GGCGCGCTTGTAAGCCTTAATTAATATGCA |
| WZH567-pxyfpn-2-Pxyl-sfGFP-PodJ1-635-R | TTAAGGCTTACAAGCGCGCCTTCGACTTCT |
| SWD700_(start)_(sfgfp)_forward | ATATGCATGGTACCTTAAGATCTCGAATGAGCAAAGGAGAAGAACTTTTCACTGG |
| SWD790_(sfgfp)_(HRSAT)_reverse | GCCCAAGGTGGCCGACCGGTGTTTGTAGAGCTCATCCATGCC |
| SWD791_(podJ_251_670)_forward | TACAAACACCGGTCGGCCACCTTGGGCGCCGTCGAGACTG |
| SWD792_(podJ_251_670)_(stop)_reverse | ACTAGTGGATCCCCCGGGCTGCAGCTTCAGCCCAAGCGCGCCTTCGAC |

### Table S3. List of plasmids constructed

| Plasmid | Description | Reference |
|---|---|---|
| pTEV5 | bacterial expression vector | Rocco 2008 |
| pTEV6 | bacterial expression vector with MBP solubilization tag | Rocco 2008 |
| pXCHYC-6 | *C. crescentus* integrating C-terminal mCherry fusion vector | Thanbickler paper |
| pBXMCS-2 | *C. crescentus* high-copy replicating C-terminal M2 fusion vector | Thanbickler paper |
| pWZ21 | pACYC-PleC-mCherry | this study |
| pWZ24 | pBAD-YFP-PodJ | this study |
| pWZ12 | pCDF-YFP-PodJ | this study |
| podj02 | pCDF-YFP-PodJ(1-702) | this study |
| podj17 | pCDF--YFP-PodJ(1-470, 643-702) | this study |
| podj23 | DH5α pCDF-YFP-PodJ(1-635) | this study |
| podj31-7 | pCDF-YFP-PodJ(1-588, 643-702) | this study |
| podj33 | pCDF-YFP-PodJ(1-470, 589-702) | this study |
| podj32 | pCDF-YFP-PodJ(1-470, PopZ 24-102, 643-702) | this study |
| pWZ138-kan | pBVMCS-6-Pvan-Plec-mCherry | this study |
| pWZ138-chlor | pBVMCS-2-Pvan-Plec-mCherry | this study |
| pCZ340 | pCDF-PleC(Atu)-CFP | this study |
| pCZ341 | pCDF-PleC(Xan)-CFP | this study |
| pCZ342 | pCDF-PleC(Hyp)-CFP | this study |
| plec2 | pBAD-PleC(1-53, 302-842)-mcherry | this study |
| plec3 | pBAD-PleC(1-301, 551-842)-mcherry | this study |
| plec4 | pBAD-PleC(1-550)-mcherry | this study |
| pWZ127 | pBAD-PleC-PAS C-mcherry | this study |
| pWZ135 | pBAD-PleC-PAS D-mcherry | this study |
| pWZ136 | pBAD-PleC-PAS CD-mcherry | this study |
| pBAD-PleC-delta PAS C | pBAD-PleC-delta PAS C-mcherry | this study |
| pBAD-PleC-delta PAS D | pBAD-PleC-delta PAS D-mcherry | this study |
| YFP-J23102-pCZ267 | pACYCDuet-J23102-YFP-PleC(302-548)-CcaS(502-753) | this study |
| pCZ259 | pProTet.E333-CcaR | this study |
| pCZ253 | pACYCDuet-CcaS(1-57)-PleC(302-548)-CcaS(502-753) | this study |
| pW173 | pBAD-YFP-PodJ(1-470, 636-974) | this study |
| pCZ412 | pBAD-YFP-PodJ(470-635) | this study |
| pWZ24-dPeri | pBAD-YFP-PodJ(1-702) | this study |
| pBAD vector | pBAD empty vector | this study |
| pCZ253(H534A) | pACYCDuet-CcaS(1-57)-PleC(302-548)-CcaS(502-753, H534A) | this study |
| pCZ259(D51A) | pProTet.E333-CcaR(D51A) | this study |
| pBVMCS-6-PleC delta CD-mCherry | pBVMCS-6-PleC delta CD-mCherry | this study |

| | | |
|---|---|---|
| pBVMCS-6-PleC delta C-mCherry | pBVMCS-6-PleC delta C-mCherry | this study |
| pBVMCS-6-PleC delta D-mCherry | pBVMCS-6-PleC delta D-mCherry | this study |
| pCZ253 (D433A, T435A) | pACYCDuet-CcaS(1-57)-PleC(302-548, D433A, T435A)-CcaS(502-753) | this study |
| pCZ253 (D548A, I549V, T550A) | pACYCDuet-CcaS(1-57)-PleC(302-548, D548A, I549V, T550A)-CcaS(502-753) | this study |
| pCZ253 (D433A, T435A, D548A, I549V, T550A) | pACYCDuet-CcaS(1-57)-PleC(302-548, D433A, T435A, D548A, I549V, T550A)-CcaS(502-753) | this study |
| pCZ391 | pTEV5-sfGFP | this study |
| pmas00001 | pTEV5-mCherry | this study |
| pCZ388 | pTEV5-sfGFP-PodJ 1-635 | this study |
| pmas00002 | pTEV5-PleC-AB-HK-mCherry | this study |
| pCZ388-dCC-13 | pTEV5-sfGFP-PodJ 250-635 | this study |
| pCZ382 | pTEV5-SNAP-PodJ 1--635 | this study |
| pCZ479 | pBAD-YFP-PodJ 250-635 | this study |
| WZ365 | pxyfpn-2-Pxyl-sfGFP-PodJ1-635 | this study |
| pSWD259 | pXyl-2-sfgfp-hrsat-podJ(251-635) | this study |

### Table S4. List of all strains used in this study

| Strain | Description | Plasmid Number | Reference |
|---|---|---|---|
| *E. coli* DH5α | bacterial cloning strain | | Invitrogen |
| *E. coli* BL21 | bacterial expression strain | | Novagen |
| BW29655 | bacterial strain for reporter gene assay | | |
| *C. crescentus* NA1000 | laboratory *Caulobacter crescentus* strain | | |
| WSC1292 | DH5α pACYC-PleC-mCherry | pWZ21 | this study |
| WSC1347 | BL21(DE3) pACYC-PleC-mCherry | pWZ21 | this study |
| WSC1234 | DH5α pBAD-YFP-PodJ | pWZ24 | this study |
| CZ668 | BL21(DE3) pBAD-YFP-PodJ | pWZ24 | this study |
| WSC1232 | DH5α pCDF-YFP-PodJ | pWZ12 | this study |
| WSC1343 | BL21(DE3) pCDF-YFP-PodJ | pWZ12 | this study |
| WSC1354 | BL21(DE3) pCDF-YFP-PodJ, pACYC-PleC-mCherry | pWZ12, pWZ21 | this study |

| | | | |
|---|---|---|---|
| WSC1237 | DH5α pCDF-YFP-PodJ(1-702) | pod02 | this study |
| WSC1367 | BL21(DE3) pCDF-YFP-PodJ(1-702) | pod02 | this study |
| WSC1258 | DH5α pCDF-YFP-PodJ(1-635) | podj23 | this study |
| WSC1252 | DH5α pCDF--YFP-PodJ(1-470, 643-702) | podj17 | this study |
| WSC1344 | BL21(DE3) pCDF--YFP-PodJ(1-470, 643-702) | podj17 | this study |
| WSC1266 | DH5α pCDF-YFP-PodJ(1-588, 643-702) | podj31-7 | this study |
| WSC1341 | BL21(DE3) pCDF-YFP-PodJ(1-588, 643-702) | podj31-7 | this study |
| WSC1348 | BL21(DE3) pCDF-YFP-PodJ(1-588, 643-702), pACYC-PleC-mCherry | pod31-7, pWZ21 | this study |
| WSC1268 | DH5α pCDF-YFP-PodJ(1-470, 589-702) | podj33 | this study |
| WSC1342 | BL21(DE3) pCDF-YFP-PodJ(1-470, 589-702) | podj33 | this study |
| WSC1351 | BL21(DE3) pCDF-YFP-PodJ(1-470, 589-702), pACYC-Plec-mcherry | pod33, pWZ21 | this study |
| WSC1267 | DH5α pCDF-YFP-PodJ(1-470, PopZ 24-102, 643-702) | podj32 | this study |

| | | | |
|---|---|---|---|
| WSC1208 | NA1000 pBVMCS-6-Pvan-Plec-mCherry | pWZ138 | this study |
| WSC1209 | DpodJ, pBVMCS-6-Pvan-Plec-mCherry | pWZ138 | this study |
| WSC1210 | Delta podJ, pBVMCS-6-Pvan-Plec-mCherry, pXYFPN-2-Pxyl-sfGFP-PodJ | pWZ138 | this study |
| WSC1221 | Delta podJ, pXYFPN-2-Pxyl-sfGFP-PodJΔ471-635, pBVMCS-6-Pvan-Plec-mCherry | pWZ138 | this study |
| CZ588 | BL21(DE3) pCDF-PleC(Atu)-CFP | pCZ340 | this study |
| CZ565 | BL21(DE3) pBAD-YFP-PodJ, pCDF-PleC(Atu)-CFP | pWZ24, pC340 | this study |
| CZ589 | BL21(DE3) pCDF-PleC(Xan)-CFP | pCZ341 | this study |
| CZ564 | BL21(DE3) pBAD-YFP-PodJ, pCDF-PleC(Xan)-CFP | pWZ24, pC341 | this study |
| CZ590 | BL21(DE3) pCDF-PleC(Hyp)-CFP | pCZ342 | this study |
| CZ563 | BL21(DE3) pBAD-YFP-PodJ, pCDF-PleC(Hyp)-CFP | pWZ24, pC342 | this study |
| WSC1297 | DH5α pBAD-PleC(1-53, 302-842)-mcherry | plec2 | this study |
| WSC1298 | DH5α pBAD-PleC(1-301, 551-842)-mcherry | plec3 | this study |

| | | | |
|---|---|---|---|
| WSC1299 | DH5α pBAD-PleC(1-550)-mcherry | plec4 | this study |
| WSC1300 | DH5α pBAD-PleC-PAS A-mcherry | pWZ127 | this study |
| WSC1301 | DH5α pBAD-PleC-PAS B-mcherry | pWZ135 | this study |
| WSC1302 | DH5α pBAD-PleC-PAS AB-mcherry | pWZ136 | this study |
| WSC1303 | DH5α pBAD-PleC-delta PAS A-mcherry | pBAD-PleC-delta PAS C | this study |
| WSC1304 | BL21(DE3) pBAD-PleC-delta PAS B-mcherry | pBAD-PleC-delta PAS D | this study |
| WSC1210 | Delta PodJ, pBVMCS-6-Pvan-PleC-mCh, pXYFPN-2-Pxyl-sfGFP-PodJ | pWZ138 | this study |
| CZ446 | BW29655, pACYCDuet-J23102-YFP-PleC(302-548)-CcaS(502-753) | YFP-J23102-pCZ267 | this study |
| CZ442 | BW29655, pACYCDuet-J23102-YFP-PleC(302-548)-CcaS(502-753), pBAD-CFP-PodJ,pProTet.E333-CcaR | YFP-J23102-pCZ267, pWZ74, pCZ259 | this study |
| CZ388 | BW29655, pACYCDuet-CcaS(1-57)-PleC(302-548)-CcaS(502-753), pBAD-YFP-PodJ,pProTet.E333-CcaR | pCZ253, pWZ24, pCZ259 | this study |

| | | | |
|---|---|---|---|
| CZ145 | BW29655, pACYCDuet-CcaS(1-57)-PleC(302-548)-CcaS(502-753), pBAD-YFP-PodJ(1-470, 636-974),pProTet.E333-CcaR | pCZ253, pW173, pCZ259 | this study |
| CZ84 | BW29655, pACYCDuet-CcaS(1-57)-PleC(302-548)-CcaS(502-753), pBAD-YFP-PodJ(470-635),pProTet.E333-CcaR | pCZ253, pCZ412, pCZ259 | this study |
| CZ639 | BW29655, pACYCDuet-CcaS(1-57)-PleC(302-548)-CcaS(502-753), pBAD-YFP-PodJ(1-702),pProTet.E333-CcaR | pCZ253, pWZ24-dPeri, pCZ259 | this study |
| CZ612 | BW29655, pACYCDuet-CcaS(1-57)-PleC(302-548)-CcaS(502-753), pBAD vector,pProTet.E333-CcaR | pCZ253, pBAD vector, pCZ259 | this study |
| CZ619 | BW29655, pACYCDuet-CcaS(1-57)-PleC(302-548, D433A, T435A)-CcaS(502-753), pBAD-YFP-PodJ,pProTet.E333-CcaR | pCZ253(D433A, T435A), pWZ24, pCZ259 | this study |
| CZ704 | BW29655, pACYCDuet-CcaS(1-57)-PleC(302-548, D548A, I549V, T550A)-CcaS(502-753), pBAD-YFP-PodJ,pProTet.E333-CcaR | pCZ253(D548A, I549V, T550A), pWZ24, pCZ259 | this study |
| CZ706 | BW29655, pACYCDuet-CcaS(1-57)-PleC(302-548, D433A, T435A, D548A, I549V, T550A)-CcaS(502-753), pBAD-YFP-PodJ,pProTet.E333-CcaR | pCZ253(D346A, D348A, D433A, D435A), pWZ24,pCZ259 | this study |

| | | | |
|---|---|---|---|
| CZ633 | BW29655, pACYCDuet-CcaS(1-57)-PleC(302-548, D433A, T435A)-CcaS(502-753), pBAD-YFP-PodJ,pProTet.E333-CcaR | pCZ253(D433A, T435A), pBAD vector, pCZ259 | this study |
| CZ633 | BW29655, pACYCDuet-CcaS(1-57)-PleC(302-548, D548A, I549V, T550A)-CcaS(502-753), pBAD-YFP-PodJ,pProTet.E333-CcaR | pCZ253(D548A, I549V, T550A), pBAD vector, pCZ259 | this study |
| CZ634 | BW29655, pACYCDuet-CcaS(1-57)-PleC(302-548, D433A, T435A, D548A, I549V, T550A)-CcaS(502-753), pBAD-YFP-PodJ,pProTet.E333-CcaR | pCZ253(D346A, D348A, D433A, D435A), pBAD vector, pCZ259 | this study |
| KAK465 | NA1000, pBVMCS-2 | pBVMCS-2 | this study |
| KAK466 | Delta PleC, pBVMCS-2 | pBVMCS-2 | this study |
| KAK343 | Delta PleC, pBVMCS-6-PleC-mCherry | pBVMCS-6-PleC-mCherry | this study |
| KAK344 | Delta PleC, pBVMCS-6-PleC delta AB-mCherry | pBVMCS-6-PleC delta CD-mCherry | this study |
| KAK345 | Delta PleC, pBVMCS-6-PleC delta A-mCherry | pBVMCS-6-PleC | this study |

| | | | |
|---|---|---|---|
| | | delta C-mCherry | |
| KAK346 | Delta PleC, pBVMCS-6-PleC delta B-mCherry | pBVMCS-6-PleC delta D-mCherry | this study |
| MJC120 | Delta PleC pBVMCS-6 | pBVMCS-6 | this study |
| MJC122 | NA1000 PleC pBVMCS-6 | pBVMCS-6 | this study |
| CZ683 | Delta PleC, pBVMCS-6-PleC -mCherry | pWZ138-kan | this study |
| CZ349 | DH5α pTEV5-sfGFP | pCZ391 | this study |
| CZ149 | Rosetta pTEV5-mCherry | pmas00001 | this study |
| CZ604 | BL21, pTEV5-sfGFP-PodJ 1-635 | pCZ388 | this study |
| CZ147 | Rosetta pTEV5-PleC-AB-HK-mCherry | pmas00001 | this study |
| CZ703 | Rosetta, pTEV5-sfGFP-PodJ 250-635 | pCZ388-dCC-13 | this study |
| CZ602 | BL21, pTEV5-SNAP-PodJ 1--635 | pCZ382 | this study |
| WSC0398 | Rosetta, pET28-His-PopZ | | this study |
| WSC1325 | BL21, PTEV5-CCKA-cc | | this study |

| | | | |
|---|---|---|---|
| WSC1326 | BL21, PTEV5-DIVL-cc | | this study |
| CZ718 | BW29655, pACYCDuet-CcaS(1-57)-PleC(302-548)-CcaS(502-753), pBAD-YFP-PodJ 250-635,pProTet.E333-CcaR | pCZ253, pCZ479, pCZ259 | this study |
| SWD655 | NA1000, podJ::chlor PpopZ::mCherry-PopZ | WZ365 | this study |
| SWD649 | NA1000, podJ::chlor | pSWD259 | this study |

**Text S1**

**Construction of plasmids**

The plasmids used in this study can be found in the Plasmids section Table S3. Below is an example of the generation of plasmid using the Gibson assembly approach based on Table S3 information. This can also be similarly applied to construct other plasmids used in this study. Oligonucleotide primers applied for amplification of the gene insert are designed using the j5 online program, and they featured overlaps of 26 bases to the insertion site in the plasmid.[10]. Oligonucleotides were synthesized by IDT (Coralville, IA), and all DNA sequencing reactions were performed by Genewiz (South Plainfield, NJ). A Gibson reaction master mix was prepared from 5x reaction buffer, T5 exonuclease (NEB), Phusion polymerase (NEB), Taq ligase (NEB), and stored as aliquots of 15 μl at -20°C[11]. The plasmid pCZ253 is designed for the expression of chimera sensor PleC-CcaS based on the backbone of pACYCDuet with the fusion of CcaS(1-57)-PleC(302-548)-CcaS(502-753) driven by a constitutive promoter. Amino acids 1-57 of CcaS are fused to 302-548 of PleC, followed by residues 502-753 from CcaS. The vector and insert were linearized through PCR using standard protocols for Phusion polymerase described in the plasmid cloning strategies section with primer pairs of CZ583/CZ524 and CZ525/CZ584, respectively. PCR products with 26 base pair overhangs for Gibson assembly were constructed following a molar ratio of linearized vector: insert = 1:10 (at least 100ng for the vector). An annealing temperature of 55°C for 1h was used, followed by 10 min at 4°C and 10 μl were then transformed into chemically competent *E. coli* DH5a cells using the KCM transformation method and plated on LB/agar supplemented with the appropriate antibiotic. Colonies were screened using primers CZ525/CZ584sing DreamTaq PCR with a 1-minute extension time.
